# The intrinsically disordered region of the E3 ubiquitin ligase TRIP12 induces the formation of chromatin condensates and interferes with DNA damage response

**DOI:** 10.1101/2023.09.13.556486

**Authors:** Claire Vargas, Manon Brunet, Alban Ricard, Damien Varry, Dorian Larrieu, Fernando Muzzopappa, Fabian Erdel, Guillaume Labrousse, Naïma Hanoun, Laetitia Ligat, Manon Farcé, Alexandre Stella, Odile Burlet-Schiltz, Henrik Laurell, Nicolas Bery, Pierre Cordelier, Marlène Dufresne, Jérôme Torrisani

## Abstract

Chromatin compaction is crucial for the faithful expression and integrity of the genome. Although largely studied, proteins and mechanisms that control the chromatin compaction are not entirely discovered. We previously showed that the nuclear HECT-type E3 ubiquitin ligase Thyroid hormone Receptor Interacting Protein 12 (TRIP12) is tightly associated to chromatin. As TRIP12 is overexpressed in several types of cancers, we explored herein the consequences of a TRIP12 overexpression on chromatin homeostasis. First, we established the TRIP12 proxisome and unveiled its pleiotropic role in chromatin regulation. Second, we demonstrated that TRIP12 overexpression leads to the formation of chromatin condensates enriched in heterochromatin marks via its intrinsically disordered region (IDR). We further discovered that the formation of TRIP12-mediated chromatin condensates is highly dynamic and driven by a mechanism of phase separation. Chromatin condensate formation depends on the TRIP12 concentration, the length of the TRIP12-IDR and relies on electrostatic interactions. We found that the formation of TRIP12 mediated-condensates alters cell cycle progression, genome accessibility, transcription as well as DNA damage response by inhibiting the accumulation of Mediator of DNA Damage Checkpoint 1 (MDC1). Altogether, this study reveals a novel dynamic role for TRIP12 in chromatin compaction independently of its ubiquitin ligase activity with important consequences on cellular homeostasis.

**GRAPHICAL ABSTRACT:** 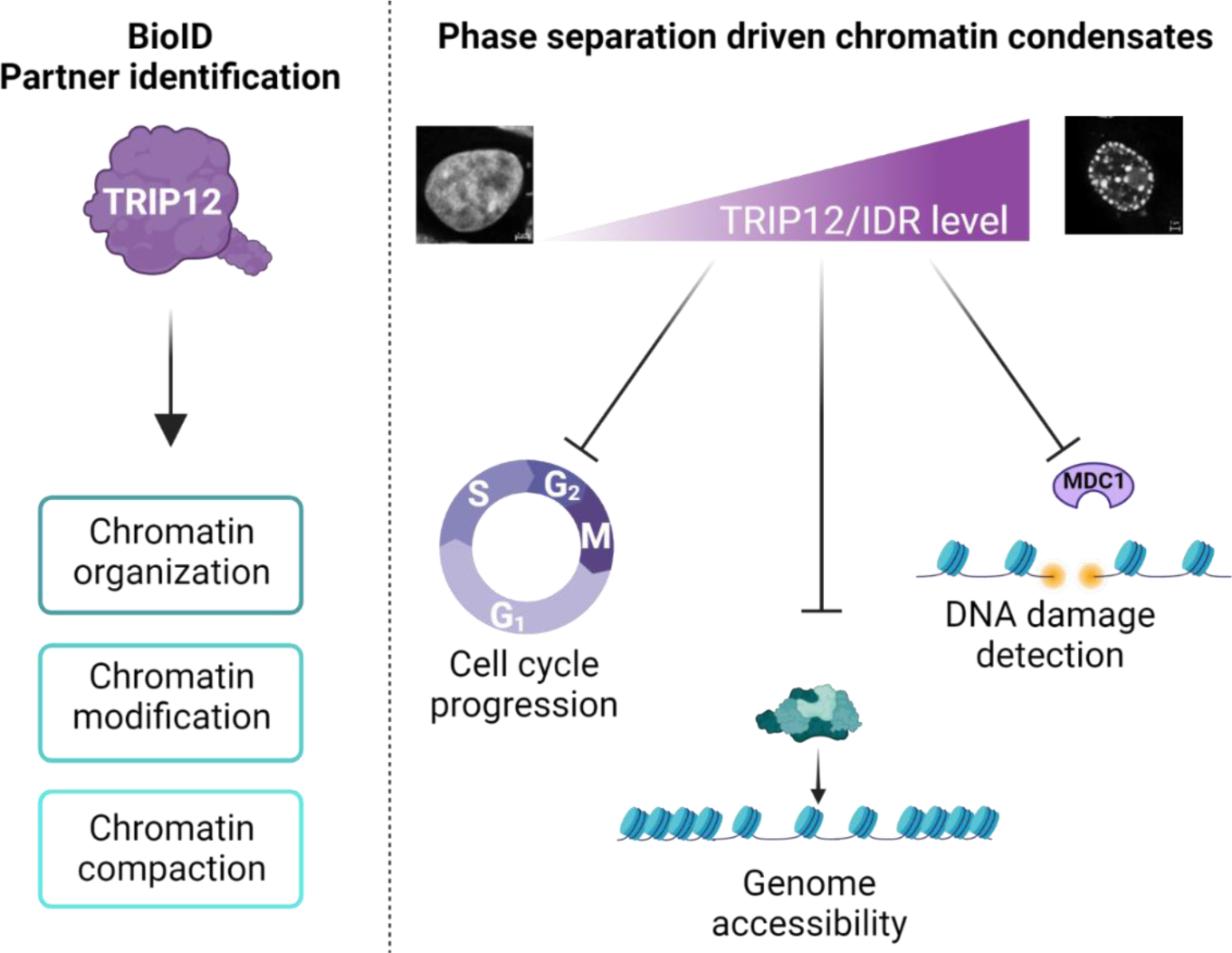

## INTRODUCTION

Chromatin is classically classified in two main categories: euchromatin and heterochromatin. Euchromatin is composed of transcriptionally active and relaxed chromatin that is modified by specific histone marks (i.e.: H3K27ac). In contrast, the more compacted heterochromatin contains fewer active genes and is enriched in a different set of histone marks (i.e.: H3K9me3 and H3K27me3). Chromatin compaction is known to condition a multitude of biological processes such as gene expression by modulating the accessibility of the genome to the transcriptional machinery. It also participates in the efficacy of the DNA damage response (DDR)^1,2^. Although largely studied, proteins and mechanisms that control the chromatin compaction are not entirely understood.

Thyroid hormone Receptor Interacting protein 12 (TRIP12) is an E3 ubiquitin ligase that belongs to the important ubiquitin-proteasome system. TRIP12 is a nuclear protein that has been involved in several biological processes such as the control of cell proliferation^3^, DDR^4,5^ and cellular differentiation^6–8^. Importantly, TRIP12 has been implicated in all of these processes via the action of its HECT (Homologous to E6-AP carboxyl-terminus) ubiquitinating catalytic domain^6,8^. For instance, we previously discovered that TRIP12 participates in pancreatic differentiation by inducing the degradation of the Pancreas Transcription Factor 1A (PTF1a)^8^. Moreover, TRIP12 regulates chromatin remodeling by controlling the stability of Brg1 associated factor 57 (BAF57), a component of the switch/sucrose non-fermenting (SWI/SNF) complex^9^. TRIP12 ensures the degradation of Additional Sex Comb Like 1 (ASXL1), a key regulator of Polycomb Repressing Complex 1 (PRC1) and it is also present in the interactome of the PRC components such as Embryonic Ectoderm Development (EED), Suppressor of Zeste 12 protein homolog (SUZ12) and Enhancer of Zeste Homologue 2 (EZH2)^10–12^. TRIP12 induces the degradation of the transcription factor Ying Yang 1 (YY1), a key component of the INO80 remodeling complex^13^. The implication of TRIP12 in chromatin homeostasis is also supported by the fact that TRIP12 is one of the most abundant interacting partners of the histone 2B (H2B)^14^.

Our previous study identified an intrinsically disordered region (IDR) at the N-terminal extremity of TRIP12 that ensures its tight interaction with chromatin^15^. IDRs are protein regions that lack tertiary structure^16^. Despite their presence in almost half of the proteome, their size varies from several residues that serve as linkers between structured domains to several hundred residues, which facilitate interactions with nucleic acids or proteins^16^. IDR-containing-proteins also participate in chromatin organization and gene expression by inducing biophysical processes, such as liquid-liquid (LLPS) or polymer-polymer (PPPS) phase separation (e.g.: BRD4 (Bromodomain-containing protein 4) and MED1 (Mediator of RNA polymerase II transcription subunit 1))^17,18,19^.

TRIP12 alterations have been involved in several pathologies. Indeed, TRIP12 gene mutations are tightly associated with intellectual disabilities such as Clark-Baraitser syndrome^20,21^ and autism spectrum disorders^22–24^. It is also reported that TRIP12 protein is overexpressed in several types of cancer and preneoplastic lesions^3,25^.

In this study, we sought to better understand the modes of action and the consequences of TRIP12 overexpression on chromatin homeostasis. We confirmed the pleiotropic role for TRIP12 in chromatin organization but more importantly, we discovered that its IDR is implicated in the formation of chromatin condensates with important consequences on DDR.

## MATERIAL AND METHODS

### Cell culture and treatments

HelaS3 (ATCC CCL-2™), HEK-293T (CRL-3216™), U2OS (HTB-96^TM^), HCT-116 (CCL-247^TM^) and H2B-dsRed HelaS3^15^ cells were grown in DMEM 4.5 g/L glucose medium and hTERT RPE-1 (CRL-4000^TM^) were grown in advanced DMEM/F-12 4.5 g/L glucose medium both supplemented with 10% fetal calf serum (FCS), L-glutamine and antibiotics (Life Technologies) at 37°C in humid atmosphere with 5% CO_2_. Cells were routinely tested for mycoplasma contamination. HelaS3 cells were treated with 10 µM 5-ethynyl 2’-deoxyuridine (EdU) (Invitrogen) for 15 min to visualize replicated DNA. The EdU incorporation was visualized using Click-It™ EdU Alexa Fluor™ 647 Imaging kit (Invitrogen). HelaS3 were treated with 1% of 1,6-Hexanediol (Tocris 7046) or vehicle for 18 h at 37°C to assess the stability of chromatin condensates. Cell lines were X-irradiated at 1 Gray with a XRAD SmART irradiator device to induce DNA damages.

### Generation of inducible GFP-degrader Hela S3 cell line

Full-length VHL (Von Hippel-Lindau) cDNA (amino acids 1-213) was cloned into *Pml* I/*Xho* I sites of the pEF empty plasmid. NbGFP4 nanobody cDNA^26^ was inserted into pEF-VHL-MCS using *Nco* I/*Not* I sites. A FLAG-tag was added by PCR at the C-terminal end of the NbGFP4 degrader plasmid. VHL-NbGFP4-FLAG sequence was subsequently cloned into TLCV2 lentivector (Addgene #87360^27^) by PCR using *Age* I/*Nhe* I sites. Lentiviral production is described in a following section. HelaS3 cells were transduced for 48 h in 6 well plate in 1 ml of medium containing 8 µg/mL of polybrene (Sigma-Aldrich). VHL-NbGFP4-FLAG HelaS3 cell line was obtained after selection with 1 µg/mL of puromycin (Invivogen).

### Plasmids generation and transient cell transfection

pcDNA. 6.2-C-EmGFP-TRIP12, pcDNA. 6.2-C-EmGFP-TRIP12-C1959A, pcDNA.6.2-C-EmGFP-ΔHECT, pcDNA. 6.2-C-EmGFP-ΔIDR, pcDNA. 6.2-C-EmGFP-IDR, pcDNA.6.2-C-EmGFP-ARM-WWE, pcDNA.6.2-C-EmGFP-ΔIDR/ARM/WWE/HECT, pcDNA 6.2-C-EmGFP-HECT, pcDNA 6.2-C-EmGFP-(1-325), pcDNA 6.2-C-EmGFP-(107-445), pcDNA 6.2-C-EmGFP-(1-208), pcDNA 6.2-C-EmGFP-(108-325), pcDNA 6.2-C-EmGFP-(208-445), pcDNA 6.2-C-EmGFP-(1-107), pcDNA 6.2-C-EmGFP-(108-207), pcDNA 6.2-C-EmGFP-(208-324), pcDNA 6.2-C-EmGFP-(325-445) constructs were already described^15^. pEGFP-C2-MDC1 and pAc-GFP-TRIP12 plasmids that encodes a 2040 aa-isoform of TRIP12 were obtained from Dr J. Lukas (Copenhagen, Denmark). Cysteine C2025 catalytic site was mutated in alanine by site directed mutagenesis as previously described^8^. pstv6-NBirA* DEST and NBirA*-FLAG-EGFP-NLS plasmids were obtained from Dr AC. Gingras (University of Toronto, Canada)^14^. pDONR-201-TRIP12 containing human TRIP12 cDNA (1992 aa isoform) was previously described^8^. TRIP12 cDNA was transferred in pstv6-NBirA*-pDest using Gateway strategy (Invitrogen) to generate pstv6-NBirA*-FLAG-TRIP12 plasmid. MED1-IDR containing plasmid was obtained from Addgene #145276 and inserted into pcDNA^TM^6.2-C-EmGFP-DEST supplemented with the Simian Virus 40 NLS sequence by Gateway strategy (Invitrogen) using primers described in Suppl. Table 1. GFP-DNMT1 plasmid was obtained from Dr M. Szyf (University McGill, Montreal, Canada). RFP-H2B and H2B-GFP plasmids were a gift from Dr D. Llères (Institute of Molecular Genetics of Montpellier, France). The plasmid pDest-RFP-C1 was obtained from Dr. T. Johansen (Tromsø, Norway). The insertion TRIP12 cDNA and its IDR into the pDest-RFP-C1 plasmid was performed using Gateway strategy (Invitrogen) to generate the pDest-RFP-TRIP12 and pDest-RFP-IDR plasmids.

The different plasmids were transiently transfected in HelaS3 or VHL-NbGFP4-FLAG HelaS3 cells using jetPRIME® transfection kit (Polyplus) following manufacturer’s recommendations. Medium was changed 4h post transfection. Cells were fixed after 24h or visualized by microscopy.

### Lentivirus production and cell transduction

Generation of lentiviral particles was adapted from the protocol described in Bery et al.^28^. Briefly, HEK-293T were transfected using jetPRIME® transfection kit (Polyplus) with 6 µg of pstv6-NBirA*-FLAG-TRIP12 or pstv6-NBirA*-FLAG-EGFP-NLS, 4 µg of psPAX2, 1.5 µg of pMD2.G and 23 µL of jetPRIME® reagent. After 48 h, supernatants were collected, centrifuged and filtered through a 0.45 µm filter (Millipore). The supernatants were concentrated using Sartorius Vivaspin™ 20 centrifugal concentrator. HelaS3 cells were seeded at a density of 10^4^ cells per well in a 48 well-dish. After 24 h, cells were incubated with 125 µL of concentrated virus in the presence of protamine sulfate (4 μg/ml) for 48 h. Transduced cells were selected using 1 µg/mL of puromycin (InvivoGen).

### Live cell microscopy

For cell cycle monitoring, cells were grown on 6-well plates (Corning) in DMEM medium without phenol red (Gibco-BRL). One µg of H2B-GFP or IDR-GFP plasmids was transfected as described above. Twenty-four hours after transfection, cells were imaged and classified in three categories (low, moderate, high) depending on their GFP fluorescence intensity determined by an IncuCyte Zoom apparatus (Sartorius). Eleven to twelve cells for each category were monitored during 30 h. For analysis of chromatin condensates dynamics, HelaS3 or VHL-NbGFP4-FLAG HelaS3 cells were grown on µ-slide 4 well (IBIDI) in DMEM medium without phenol red (Gibco-BRL). 0,25 µg of IDR-GFP plasmid was transfected as described above. One hour before image acquisition, 500 nM of SiR-DNA (SC007, Spirochrome) were added to the medium to visualize DNA. For analysis of chromatin condensates reversibility, 1 µg/mL of doxycycline was added to induce VHL-NbGFP4-FLAG degrader expression. Images were acquired every hour (5 fields, 1× Air hNA 0.75) using an Operetta® CLS high content analysis system (Perkin-Elmer). Fluorescent intensity was quantified using Harmony software (version 4.9) and FIJI software.

### Immunofluorescence and DNA granularity quantification

Cells were grown on cover slips then fixed using with 4% formaldehyde and permeabilized with 0.125% Triton X-100™. Cover slips were saturated using 5% BSA-PBS for 1 h and incubated with primary antibodies (Suppl. Table 2) overnight at 4°C. After three washes, cells were incubated with appropriate secondary antibodies Alexa Fluor®-488 anti-goat, Alexa Fluor®-555 anti-mouse/rabbit or Alexa Fluor®-647 anti-rat antibodies (Suppl. Table 2) for 2 h at room temperature. Nuclei were counterstained with 1 μM DAPI for 5 min at room temperature. Cover slips were mounted on glass slides using Fluorescent Mounting Medium (DAKO). Fluorescence was visualized using LSM 780 or 880 Fast Airyscan confocal microscope (Zeiss) with a 63× NA 1.4 oil-immersion objective and analyzed using ZEN 3.5 software (Zeiss). For DAPI granularity quantification, images of individual cells were acquired using a LSM880 confocal microscope (Zeiss) (500 × 500 pixels). Each nucleus was manually defined as a region of interested (ROI). The granularity was defined by a coefficient of variation (CV) obtained by dividing the mean of all pixel integrated intensity of the ROI by the standard deviation as previously described^29^. For simplification, the slope value was defined as the leading coefficient of the linear regression curve multiplied by 10^7^. The determination of cell cycle phase by immunofluorescence using CYCLIN A, EdU staining was previously described^15^.

### Subcellular fractionation and Western blot analysis

Cells were lysed in 100 mM Tris-HCl (pH 7.5), 1,5 mM MgCl_2_, 5 mM KCl, 5 mM DTT, 0,5% NP-40®, 0,5 mM PMSF and 10 μl/mL protease inhibitors (Sigma-Aldrich). After 10 min-incubation on ice, samples were centrifuged. Supernatants containing cytosolic proteins were discarded. Nuclear pellets were incubated in a hypertonic buffer containing 200 mM Tris-HCl (pH7,5), 0.25% glycerol, 1,5 mM MgCl_2_, 0,5 mM PMSF, 0.2 mM EDTA, 0.5 mM DTT, 0.4 M NaCl and 10 μl/mL protease inhibitors. After 15 min-incubation on ice, samples were centrifuged. Supernatants containing nuclear proteins were collected. For total protein extraction, cells were lysed in RIPA buffer (50 mM Tris-HCl (pH 8), 150 mM NaCl, 0.1% SDS, 0.5% sodium deoxycholate, 1% Nonidet P-40 and 10 µl/mL protease inhibitors) for 20 min on ice. Cell lysates were sonicated 3 × 5 sec at 50% intensity (Vibra-CellTM) and centrifuged at 12 000 rpm. Proteins were denaturated in Laemmli buffer at 95°C for 5 min. Proteins were separated by 10% SDS-PAGE and transferred onto a nitrocellulose membrane (BioRad) using TransBlot Turbo apparatus. After membrane saturation and incubation with primary/secondary antibodies (Suppl. Table 2), protein expression was detected using Clarity^TM^ Western ECL Substrate (BioRad) and Chemi-DocTM XRS^+^ BioRad apparatus.

### FRAP and half-FRAP analysis

Fluorescence Recovery After Photobleaching (FRAP) and half-FRAP experiments were conducted using a confocal LSM 780 microscope (Zeiss). Bleaching was performed using the 488 nm laser at 100% power and images were collected every two seconds. Fluorescence recovery intensity was measured using ZEN black Module FRAP Efficiency Analysis. The half-FRAP analysis was performed as previously described^30^.

### BioID streptavidin pull-down purification

pstv6-NBirA*-FLAG-TRIP12 HelaS3 stable cell line was generated as described below and validated by Western blot, RT-qPCR and immunofluorescence after the addition of doxycycline (1 µg/mL). pstv6-NBirA*-FLAG-TRIP12 cells were incubated with 1 µg/mL of doxycycline and 50 µM of biotin (Sigma-Aldrich) for 24 h. During the last 15 h, cells were treated with 1.25 µM of a proteasome inhibitor MG-132 (Sigma-Aldrich). After four PBS washes, cells were scrapped in 100 mM Tris-HCl (pH 7.5), 1,5 mM MgCl_2_, 5 mM KCl, 5 mM DTT, 0,5% NP-40®, 0,5 mM PMSF and 10 μl/mL protease inhibitors (Sigma-Aldrich). After 10 min-incubation on ice, samples were centrifuged (15 min at 2.000 rpm, 4°C). Supernatants containing cytosolic proteins were discarded. Nuclear pellets were lysed with a modified RIPA (50 mM Tris-HCl (pH 8), 150 mM NaCl, 0.1% SDS, 0.5 % sodium deoxycholate, 1 % Triton X-100™ and 10 µL/mL protease inhibitors). Nuclear lysates were sonicated (5 × 6 sec at 25% intensity) using Vibra-Cell^TM^ sonicator and centrifuged (20 min at 10.000 rpm, 4°C). Clarified lysates were incubated with 40 µl of Thermo Scientific™ Pierce™ Streptavidin Magnetic Beads overnight at 4°C. Beads were washed with different buffers (1 M KCl, 0.1 M Na_2_CO_3_, 2M urea, modified RIPA) and eluted in 5% SDS, 10% glycerol, 80 mM Tris-HCl (pH 6.8), 20 mM DTT and 2 mM biotin at 95°C for 3 min. Four independent biotin pull down experiments were performed.

### Label free mass spectrometry identification (LC MS/MS)

Dried protein extracts were solubilized with 25 µl of 5% SDS. Proteins were submitted to reduction and alkylation of cysteine residues by addition of Tris(2-carboxyethyl)phosphine and chloroacetamide to a final concentration of 10 mM and 40 mM, respectively. Protein samples were processed for trypsin digestion on S-trap Micro devices (Protifi) according to manufacturer’s protocol with the following modifications: precipitation was performed using 216 µl S-Trap buffer; 1 µg Trypsin was added per sample for digestion, in 25 µl of triethylammonium bicarbonate 50 mM (pH 8,0). Tryptic peptides were resuspended in 17 µl of 2% acetonitrile and 0.05% trifluoroacetic acid and analyzed by nano-liquid chromatography (LC) coupled to tandem MS, using an UltiMate 3000 system (NCS-3500RS Nano/Cap System; Thermo Fisher Scientific) coupled to an Orbitrap QExactive Plus mass spectrometer (Thermo Fisher Scientific). Five µl of each sample were loaded on a C18 precolumn (300 µm inner diameter × 5 mm, Thermo Fisher Scientific) in a solvent composed of 2% acetonitrile and 0.05% trifluoroacetic acid, at a flow rate of 20 µl/min. After 5 min of desalting, the precolumn was switched online with the analytical C18 column (75 μm inner diameter × 50 cm, in-house packed with Reprosil C18) equilibrated in 95% solvent A (5% acetonitrile, 0.2% formic acid) and 5% solvent B (80% acetonitrile, 0.2% formic acid). Peptides were eluted using a 5%-50% gradient of solvent B over 115 min at a flow rate of 300 nl/min. The mass spectrometer was operated in data-dependent acquisition mode with the Xcalibur software. MS survey scans were acquired with a resolution of 70,000 and an AGC target of 3^e6^. The 10 most intense ions were selected for fragmentation by high-energy collision induced dissociation and the resulting fragments were analysed at a resolution of 17500 using an AGC target of 1^e5^ and a maximum fill time of 50 ms. Dynamic exclusion was used within 30 s to prevent repetitive selection of the same peptide.

### Data processing protocol

Raw MS files were processed with the Mascot software (version 2.7.0) for database search and Proline ^31^ for label-free quantitative analysis (version 2.1.2). Data were searched against Human entries of the UniProtKB protein database (release Swiss-Prot 20201022, 20,385 entries). Carbamidomethylation of cysteines was set as a fixed modification whereas oxidation of methionine was set as variable modifications. Specificity of trypsin/P digestion was set for cleavage after K or R, and two missed trypsin cleavage sites were allowed. The mass tolerance was set to 10 ppm for the precursor and to 20 mmu in tandem MS mode. Minimum peptide length was set to 7 amino acids and identification results were further validated in Proline by the target decoy approach using a reverse database at both a PSM and protein false-discovery rate of 1%. For label-free relative quantification of the proteins across biological replicates and conditions, cross-assignment of peptide ions peaks was enabled inside group with a match time window of 1 min after alignment of the runs with a tolerance of +/-600s. The list of proteins identified was compared with the list of contaminants (CRAPome) commonly found in similar approaches^32^. The mass spectrometry proteomics data were deposited to the ProteomeXchange Consortium via the PRIDE partner repository with the dataset identifier PXD045224.

### Chromatin accessibility by ATAC-seq

Genomic accessible regions were determined by two ATAC (assay for transposase accessible chromatin)-seq experiments performed by the epigenetic services of Active Motif. Briefly, HelaS3 cells were transfected with IDR-GFP construct using using jetPRIME® transfection reagent (Polyplus). After 24 h, GFP-expressing cells were sorting by cytometry using FACS Melody apparatus (BD Biosciences). 10^5^ GFP-highly positive and GFP-negative (used as controls) cells were collected. Samples were frozen accordingly to the manufacturer’s instructions in an ice-cold cryopreservation solution and stored at - 80°C until shipping. ATAC-seq reads were generated using Illumina sequencing (NextSeq 500) and were mapped to the hg38 human reference genome using BWA algorithm. MACS peak calling algorithm was used to identified the regions with high levels of tagging events. Only merged accessible regions with Max tag >50 were considered. Data are available upon request.

### RNA expression by RNA-seq analysis

Total RNA was isolated from HelaS3 cells transfected or not with IDR-GFP plasmid that were used for ATAC-seq experiments using TRIzol® Reagent (Life Technologies) according to supplier’s recommendations. Quality and quantity of total mRNA was ensured by Agilent 5400 Fragment Analyzer system. Paired-end (PE150) sequences were generated using a NovaSeq 6000 apparatus (Illumina) by Novogen (Cambridge, UK). Paired reads were aligned and mapped on *Homo sapiens* reference genome (GRCh38.p14) using STAR software. Data analysis with normalization was performed using the R software, Bioconductor packages including DESeq2 and SARTools R^33^. Sequences corresponding to IDR-GFP mRNA were discarded from the analysis. Experiments were performed in duplicate. A p-value adjustment was performed to consider multiple testing and control the false positive rate. A BH p-value adjustment was performed and the level of controlled false positive rate was set to 0.05. Data are available on GEO Omnibus data depository (GSE242544).

### UV-laser micro-irradiation

HelaS3 cells were transiently transfected with appropriate plasmids as described above. After 24 h, cells were photo-sensitized with Hoechst (Life technologies) (0.25 µg/mL) for 10 min in phenol red-free complete DMEM medium (Gibco-BRL). UV-irradiation (405 nm) was performed using FRAP module of LSM 780 Zeiss confocal microscope (10 iterations, power 100%, Plan-Apo 40×/1.3 NA) in a 37°C atmosphere with 5% CO_2_. Images were acquired every 30 sec for 60 cycles. Irradiation was performed after the 3 or 4 first acquisitions. The quantification of GFP-MDC1 recruitment on irradiated zones was done using FIJI software following procedures described in^34^.

### Bioinformatics analyses

Gene ontology enrichment and protein function networks were established using Gene Ontology (http://geneontology.org), Metascape (https://metascape.org) and String (https://string-db.org) software. Isoelectric point (pI) of TRIP12 fragments inserted in pcDNA6.2 EmGFP vectors was calculated using Expasy-Compute pI/MW tool (GFP protein sequence was omitted for the calculus). Natural disordered regions were predicted using PONDR VSL2 website (http://pondr.com). Prediction of nucleolar localization signals was determined using Nucleolar localization sequence Detector software (https://www.compbio.dundee.ac.uk/www-nod).

### Statistical analyses

For correlation determination, a Spearman r coefficient test (two tailed p value) was used with GraphPad Prism 9.5.0 software. For MDC1 and γH2AX foci number comparison, one way-ANOVA statistical test followed by multiple comparisons was used. For granularity comparison in presence or not of 1,6-hexanediol, an unpaired t test was used. *, **, *** and **** correspond to a p value <0.05, 0.01, 0.005 and 0.001, respectively.

## RESULTS

### TRIP12 is implicated in global chromatin organization and compaction networks

The TRIP12 ubiquitin ligase has been involved in numerous nuclear functions^36^ and retrieved as a prey in a number of BioGrid interactomes, in particular of two chromatin remodeling complexes (SWI/SNF and PRC2)^9,11,12^. However, a protein partner network of TRIP12 used as a bait has never been established. To fill this gap, we established a TRIP12 proxisome by biotinylation-based identification (BioID) that identifies partners at the vicinity (∼15 nm) of a protein of interest. TRIP12 was N-terminally fused to the biotin ligase NBirA*(R118G)-FLAG protein (Fig. 1A, top panel) and stably expressed in HelaS3 cells after doxycycline addition (Fig. 1A and 1B). We verified the nuclear localization of the fusion protein and the incorporation of biotin in response to doxycycline (Fig. 1A, bottom left panel). More importantly, we ensured a moderate expression of NBirA*-FLAG-TRIP12 relatively to the endogenous TRIP12 protein to limit non-specific proteins enrichment (Fig. 1B and Suppl. Fig. 1A). We also verified a chromatin-bound localization of the fusion protein on metaphasic chromosomes (Fig. 1A, right panel). After streptavidin-beads pull-down (Suppl. Fig. 1B), the identification and abundance of purified proteins were determined by mass spectrometry-based label-free quantitative proteomics. A total of 328 proteins statistically enriched at the vicinity of NBirA*-FLAG-TRIP12 were identified (Fig. 1C + Suppl. Table 3). A subset of them was randomly selected and validated by Western blot analysis (Suppl. Fig. 1C). Among the identified proteins, 18 were in common with TRIP12 BioGrid interactors such as EZH2 (Enhancer of Zest Homologue 2), SWI/SNF-Related Matrix-Associated Actin-Dependent Regulator Of Chromatin Subfamily E Member 1 (SMARCE1) or Bromodomain Containing 4 (BRD4) (Suppl. Fig. 1D). A gene ontology analysis using Metascape revealed that the most statistically enriched gene categories are “chromatin organization” and “histone modification” (Fig. 1D). In parallel, this analysis identified several functional networks among which the most statistically relevant ones correspond to chromatin organization, histone post-translational modifications, RNA splicing, epigenetic regulation and chromatin remodeling (Fig. 1E). In addition, a String analysis confirmed the close proximity of TRIP12 to proteins belonging to PRC and SWI/SNF complexes but also to other remodeling complexes such as Spt-Ada-Gcn5 acetyltransferase (SAGA), Nucleosome Remodeling and Deacetylase (NURD) and mixed lineage leukemia protein-1 (MLL1) (Fig. 1F). Of note, the three isoforms (α, β, γ) of heterochromatin protein 1 (HP1) were present in the TRIP12 proxisome (referenced as CBX5, CBX1 and CBX3 in Suppl. Table 3, respectively). Altogether, our results confirmed the tight implication of TRIP12 in global chromatin reorganization and compaction networks.

**Figure 1:**
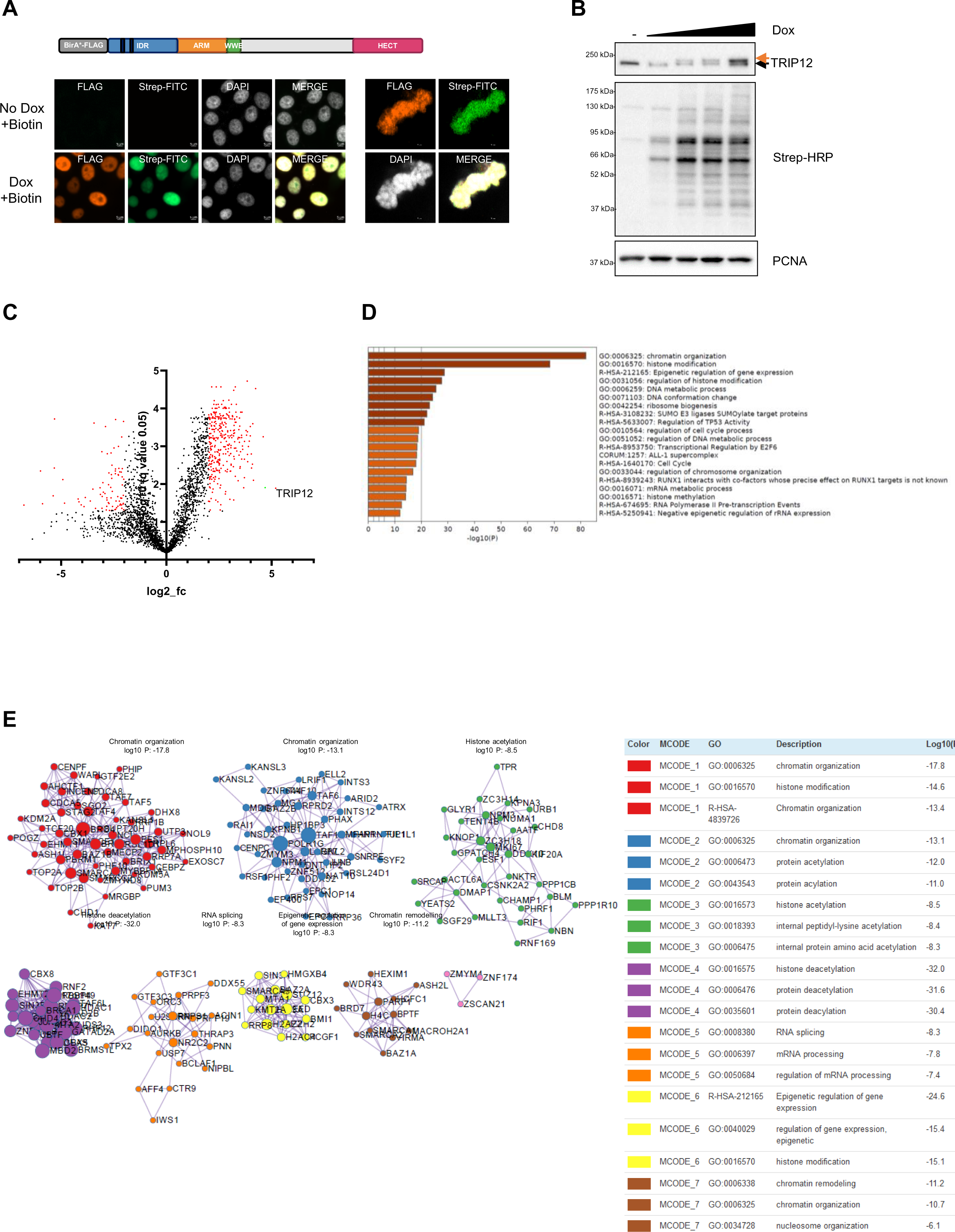
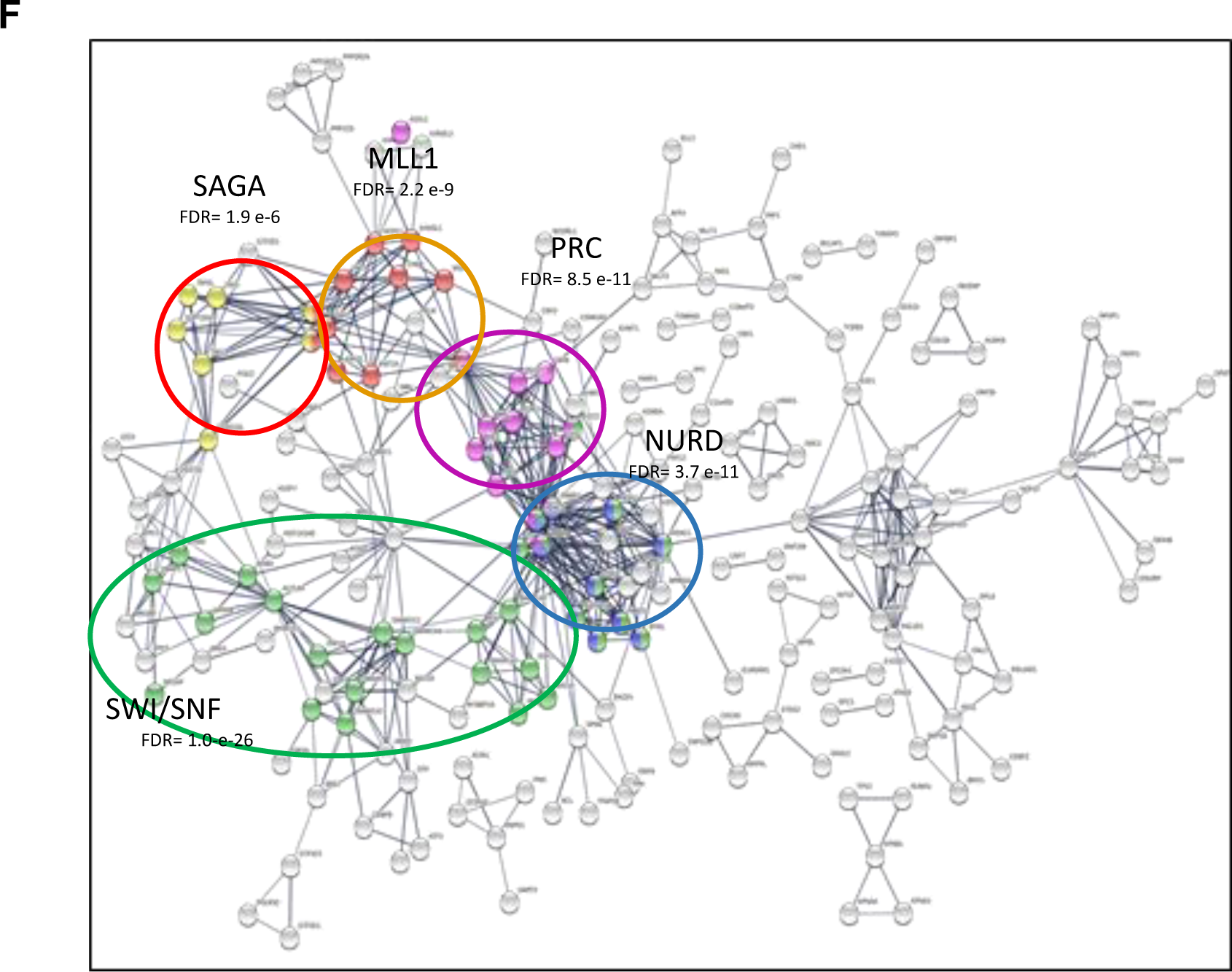
TRIP12 is implicated in global chromatin organization and compaction networks. **A-** Graphical representation of NBirA*-FLAG-TRIP12 fusion protein (top left). IDR: Intrinsically Disordered Region, ARM: armadillo repeats, WWE: tryptophan-tryptophan-glutamate rich domain and HECT: Homologous to E6-AP Carboxyl Terminus. Black rectangles represent Nuclear Localization Signals (NLS). Representative image of NBirA*-FLAG-TRIP12 and biotinylated protein localization in NBirA*-FLAG-TRIP12 expressing HelaS3 cells in interphase (bottom left) or in mitosis (bottom right) treated or not with doxycycline (1 µg/ml) and biotin (50 µM) for 24 h by immunofluorescence using an anti-FLAG antibody and streptavidin-FITC conjugate. Nuclei were counterstained with DAPI. **B-** Nuclear TRIP12 expression and biotinylation profile in response to doxycycline treatment in NBirA*-FLAG-TRIP12 expressing HelaS3 cells was determined by Western blot using TRIP12 antibody and streptavidin-HRP conjugate. Cells were treated with increasing doses of doxycycline (100, 350, 600 and 1000 ng/ml), biotin (50 µM) for 24 h and with MG-132 (1.25 µM) for 15 h. PCNA level was used as loading control. The black and orange arrows indicate endogenous and exogenous TRIP12 protein, respectively. **C-** Volcano plot of identified proteins at proximity (∼15 nm) of TRIP12 determined by BioID and label-free mass spectrometry-based quantitative proteomics. The volcano plot represents the statistical significance distribution against the log2-transformed fold change between doxycycline addition (NBirA*-FLAG-TRIP12 expression) and no doxycycline addition (Control). Proteins (n=328) were considered significantly enriched (red on the right) when displaying a corrected p value ≤ 0.05 and an absolute fold change ≥ 2. calculated from four independent experiments. Biotinylated TRIP12 protein is indicated in green on the volcano plot. **D-** Gene ontology analysis of TRIP12 interactors identified by BioID approach using Metascape software. GO terms having the highest significance are shown. **E-** Protein networks analysis using Metascape software of TRIP12 protein partners identified by BioID (left). Statistic values are expressed as log10(p-value) (right). **F-** Protein functional network using String software of TRIP12 protein partners identified by BioID. Coloured circles correspond to proteins that belong to SAGA (yellow), MLL1 (orange), PRC (violet), NURD (blue), SWI/SNF (green) complexes. Corresponding False Discovery Rate (FDR) are indicated.

### TRIP12 overexpression modifies the partitioning of chromatin independently of its catalytic activity

Proxisome identification confirmed a pleiotropic implication of TRIP12 in chromatin organization. Interestingly, a microscopic observation of the cell line used for the BioID identification revealed a clear modification of the DAPI distribution in response to NBirA*-FLAG-TRIP12 expression. Exogenous TRIP12 co-localized with large chromatin condensates and perinucleolar heterochromatin regions (Fig. 2A, left panel). These modifications were not observed after NBirA*-FLAG-EGFP-NLS expression (Fig. 2A, right panel). Therefore, we decided to further investigate the consequences of TRIP12 overexpression on global chromatin organization. We expressed different TRIP12-GFP protein constructs in HelaS3 cells (Fig. 2B). Due to the heterogeneity of transient plasmid transfection, we obtained cell populations expressing different levels of GFP fusions (Fig. 2C). In cells with a low expression level, nuclear TRIP12 was enriched in perinucleolar heterochromatin regions (Fig. 2C, left panel). Interestingly, in cells with a moderate expression level, an enlargement of perinucleolar and perinuclear heterochromatin regions concomitantly with a formation of large isolated chromatin condensates was observed (Fig. 2C, left panel). In cells with a high expression level, the nucleus was filled with even larger chromatin condensates. A quantitative analysis demonstrated a statistical positive correlation between the level of TRIP12-GFP expression and the DAPI granularity (slope: 45) that reflects the capacity of GFP constructs to modify chromatin (Fig. 2D, left graph). Of note, the formation of TRIP12-mediated chromatin condensates was neither dependent of the position nor of the GFP nor the isoform of TRIP12, as the 2040 aa-TRIP12 isoform^36^ with the GFP at the N-terminal extremity led to similar chromatin modifications (slope: 60) (Fig. 2C, right panel and Fig. 2D, right graph). Importantly, the modification of chromatin repartitioning was also observed with catalytic defective mutants (catalytic site-point mutation (C1959A)^8^ or HECT-deleted constructs), indicating that TRIP12 expression modified the spatial organization of chromatin without its ubiquitin ligase activity (Fig. 2E and 2F). Such a chromatin reorganization was not observed with well-known chromatin proteins such as the histone H2B or the large DNA methyltransferase 1 (DNMT1) (1620 aa) (Fig. 2E and 2F).

**Figure 2:**
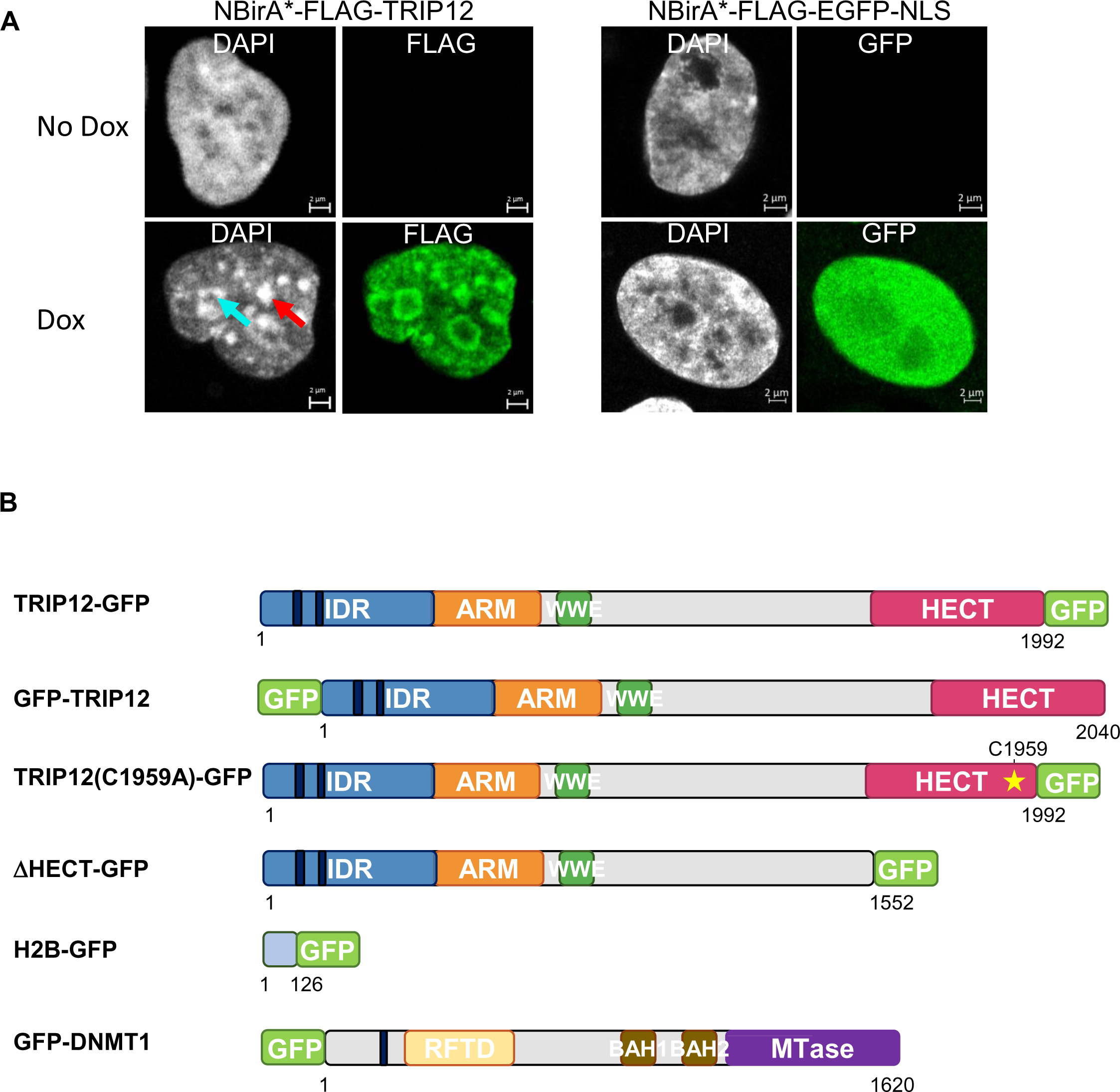
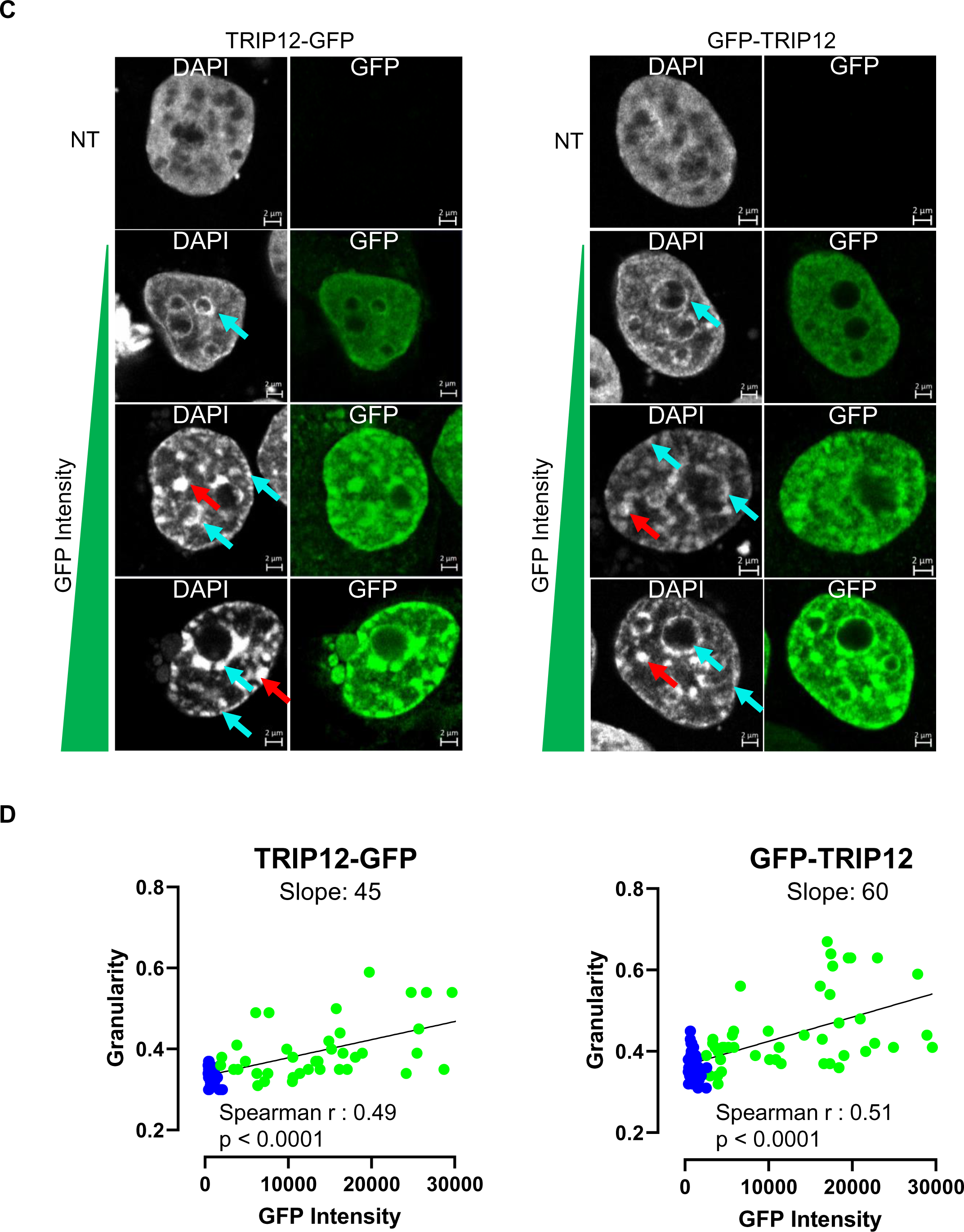
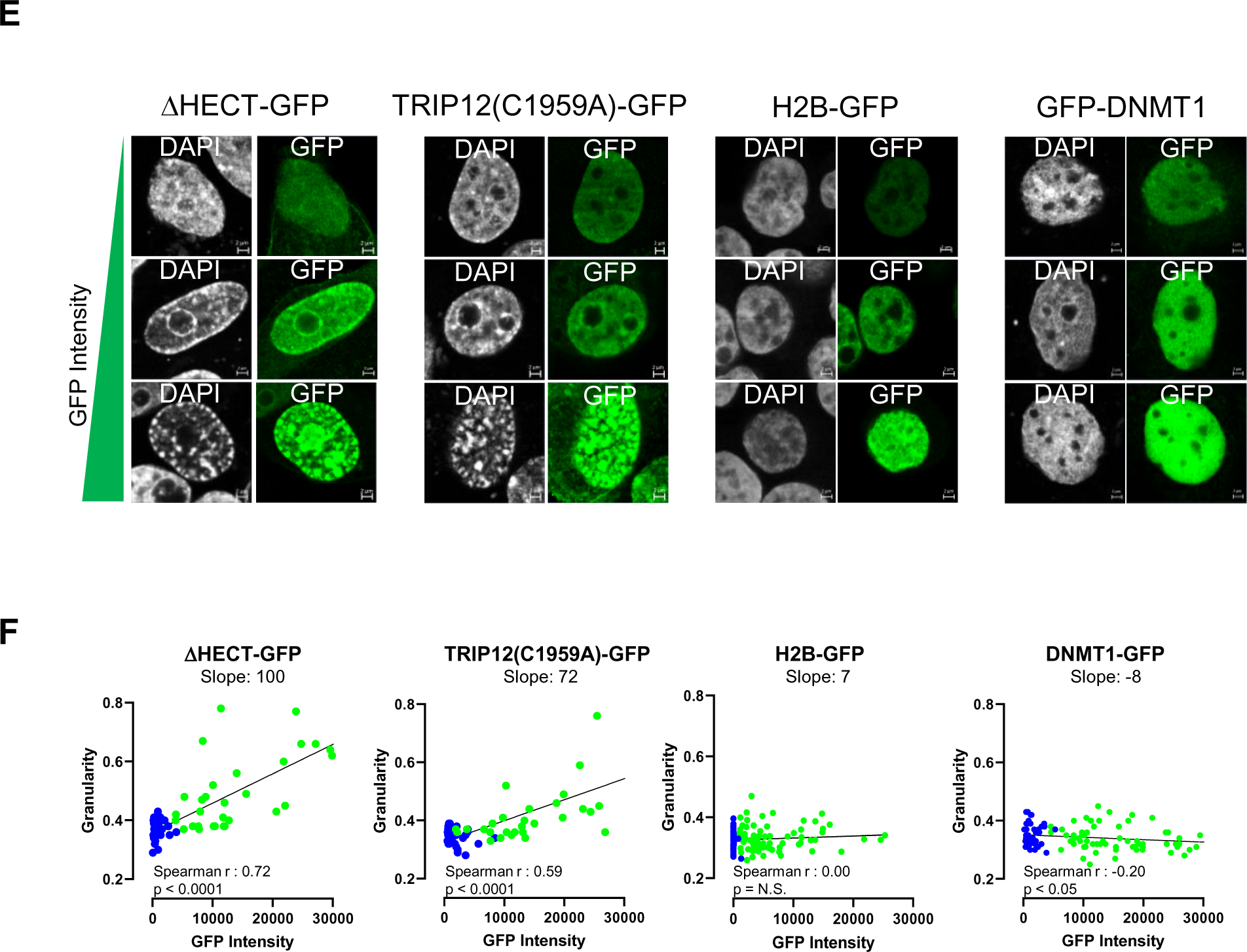
TRIP12 overexpression modifies the partitioning of chromatin independently of its catalytic activity. **A-** Representative images of NBirA*-FLAG-TRIP12 and NBirA*-FLAG-EGFP-NLS expressing HelaS3 cells treated or not with doxycycline (1 µg/ml) for 48 h by immunofluorescence using indicated antibodies. Nuclei were counterstained with DAPI. A blue arrow represents the thickening of perinucleolar regions while a red arrow represents chromatin condensates. **B-** Graphical representation of GFP fusion proteins. The yellow star locates the ubiquitin ligase catalytic site. IDR: Intrinsically Disordered Region, ARM: armadillo repeats, WWE: tryptophan-tryptophan-glutamate rich domain, HECT: Homologous to E6-AP Carboxyl Terminus, GFP: Green Fluorescent Protein, RFTD: Replication Foci Targeting Domain, BAH: Bromo-Adjacent Homology and MTase: MethylTransferase. Black rectangles represent Nuclear Localization Signals (NLS). Numbers indicate the length in amino acids of the proteins. **C-** Representative images of TRIP12-GFP (left) and GFP-TRIP12 (right) overexpressing HelaS3 cells by immunofluorescence after transient transfection. Nuclei were counterstained with DAPI. Cells with three different GFP intensities are represented. Blue arrows represent the thickening of perinuclear and perinucleolar regions. Red arrows represent chromatin condensates. **D-** Determination of DNA granularity in function to GFP expression level in TRIP12-GFP and GFP-TRIP12 expressing HelaS3 cells. For each cell, the DAPI granularity and GFP expression were determined as described in Materials and Methods on over than 50 cells. Blue and green filled circles correspond to individual non transfected and transfected cells, respectively. The linear regression curve is indicated in black. **E-** Representative images of ΔHECT-GFP, TRIP12(C1959A)-GFP, H2B-GFP and GFP-DNMT1 expressing HelaS3 cells by immunofluorescence after transient transfection. Nuclei were counterstained with DAPI. Cells with three different GFP intensities are represented. **F-** Determination of DNA granularity in function of GFP expression level in ΔHECT-GFP, TRIP12(C1959A)-GFP, H2B-GFP and GFP-DNMT1 expressing HelaS3 cells. For each cell, DAPI granularity and GFP expression were determined as described in Materials and Methods on over than 50 cells. Blue and green filled circles correspond to individual non transfected and transfected cells, respectively. The linear regression curve is indicated in black.

Taken together, our results showed that TRIP12 overexpression induces the formation of chromatin condensates in a concentration-dependent manner and independently of its catalytic activity.

### The intrinsically disordered region of TRIP12 is responsible for the formation of chromatin condensates

TRIP12 is a large protein composed of several structured domains (ARM, WWE and HECT) and a long intrinsically disordered domain (IDR)^36^ (Fig. 3A). To identify the domain(s) implicated in the formation of chromatin condensates, different fragments of TRIP12 were fused to GFP (Fig. 3B). To allow a nuclear localization, an artificial SV40 nuclear localization signal (NLS) was inserted in these constructs with the exception of the IDR-GFP construct that contains an endogenous NLS. Unlike the other domains, only the expression of IDR-GFP dramatically modified the chromatin repartitioning in a dose-dependent manner (Fig. 3C and 3D). This conclusion was further supported by the use of a TRIP12-GFP construct devoid of the IDR (ΔIDR-GFP) that failed to form chromatin condensates. This chromatin reorganization in response to IDR-GFP expression was also observed in other cell line models such as U2OS, hTert-RPE1 or HCT-116 cells (Fig. 3E).

**Figure 3:**
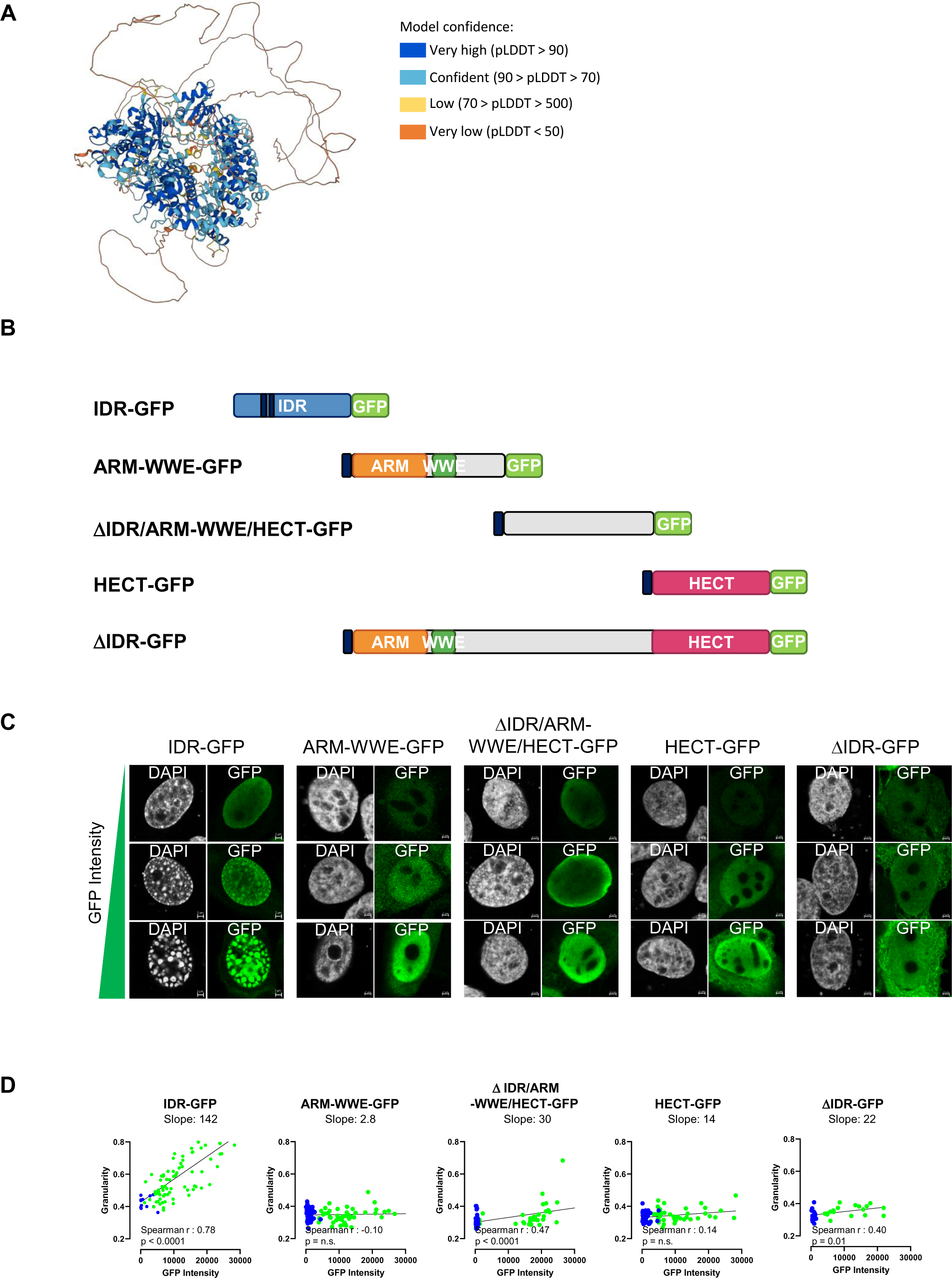
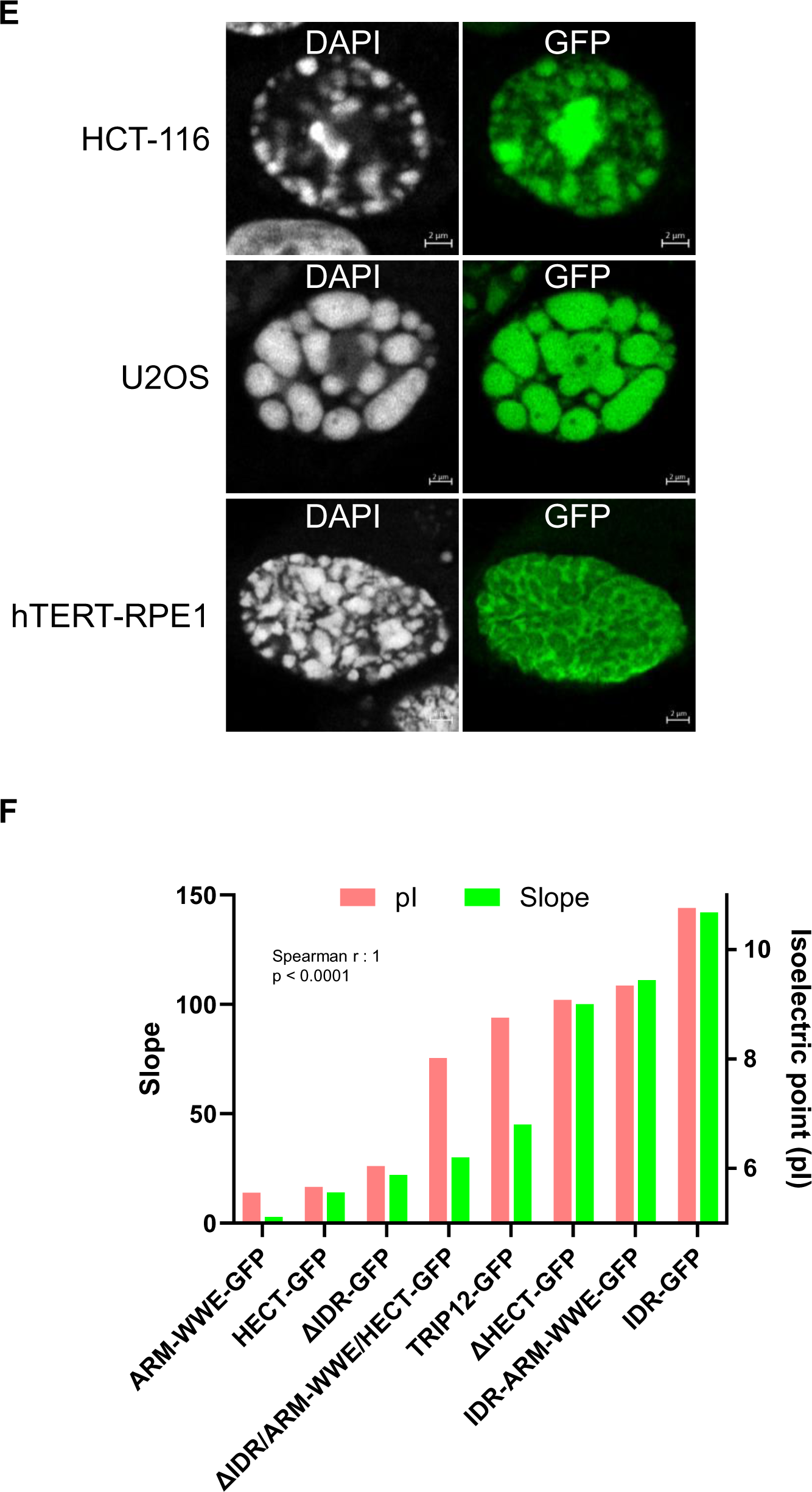
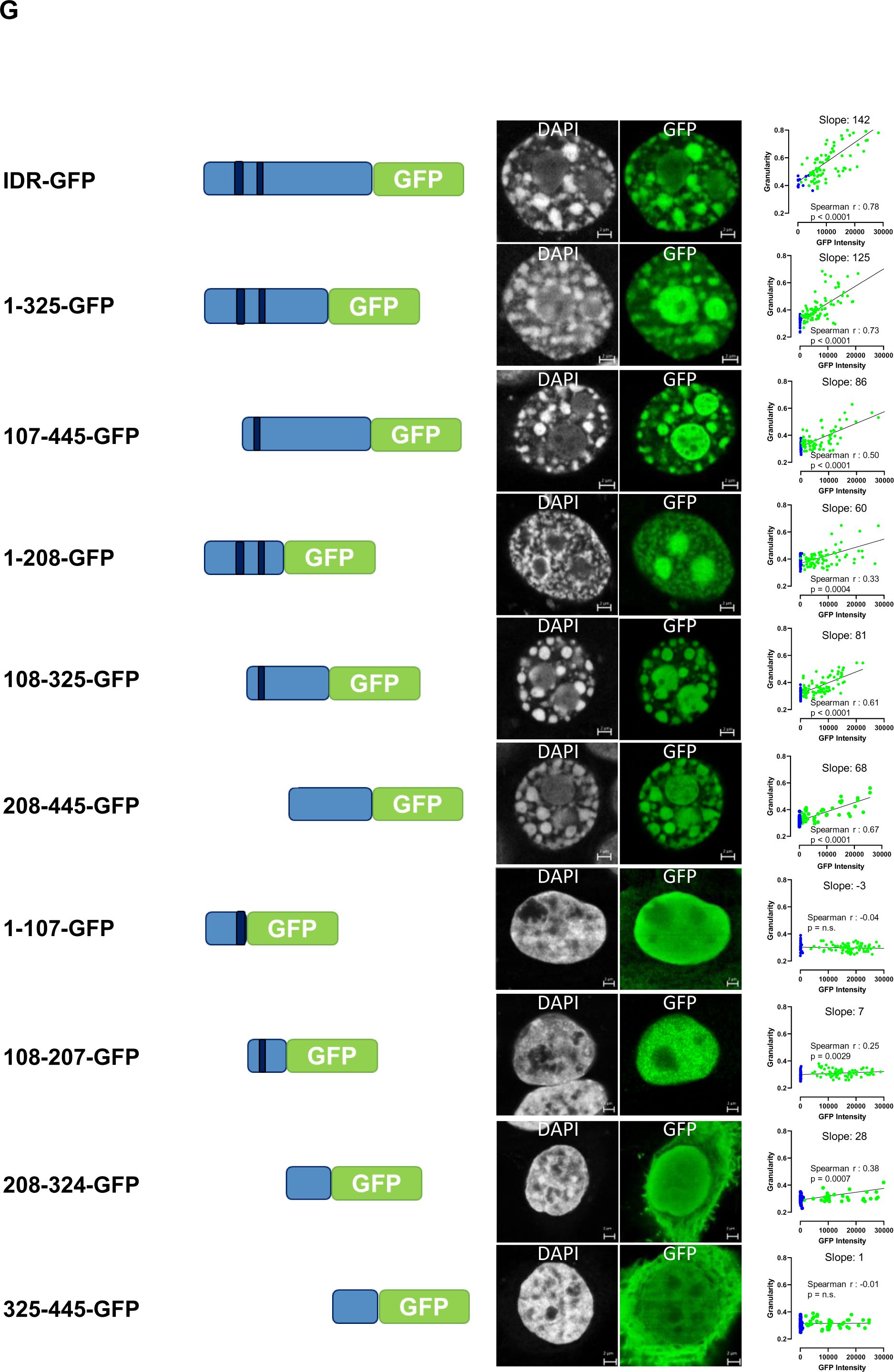
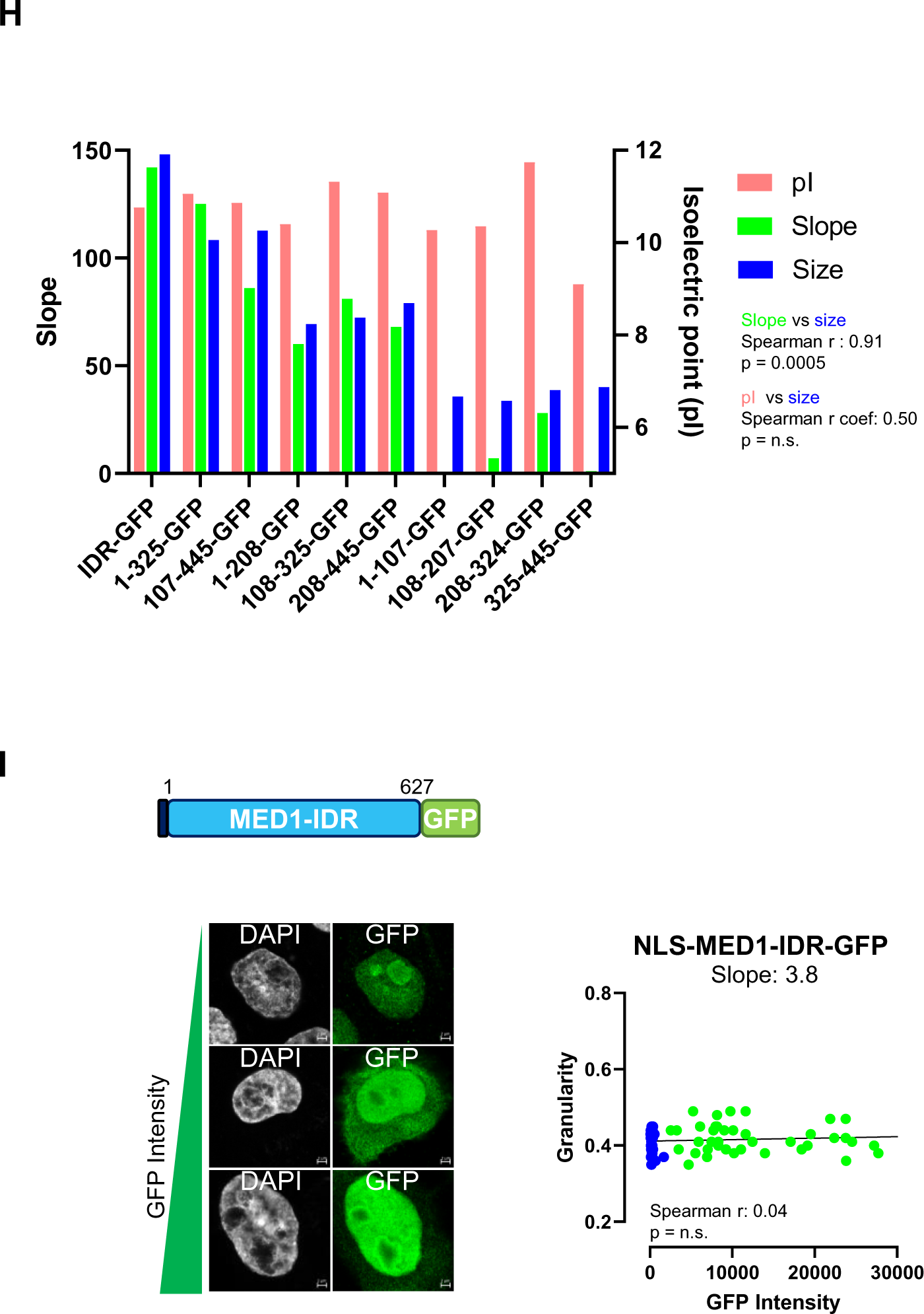
The intrinsically disordered region of TRIP12 is responsible for the formation of chromatin condensates. **A-** Prediction of TRIP12 3D-structure by AlphaFold website. Predicted local distance difference test (pLDDT) indicates the confidence level of the predicted structure. Dark blue, light blue, orange and yellow represent very high, confident, low and very low model confidence, respectively. **B-** Graphical representation of TRIP12 different domains fused to the GFP reporter protein. Dark blue rectangles locate endogenous or artificial nuclear localization signals (NLS). IDR: Intrinsically Disordered Region, ARM: armadillo domain, WWE: tryptophan-tryptophan-glutamate rich domain, HECT: Homologous to E6-AP Carboxyl Terminus and GFP: Green Fluorescent Protein. **C-** Representative images of TRIP12-domains fused to GFP expressing HelaS3 cells by immunofluorescence. Nuclei were counterstained with DAPI. Cells with three different GFP intensities are represented. Insertion of an artificial NLS in ARM-WWE-GFP, ΔIDR/ARM-WWE/HECT-GFP, HECT-GFP and ΔIDR-GFP constructs allowed a nuclear localization with a substantial leaking in the cytoplasm. **D-** Determination of DNA granularity in function of GFP expression level in TRIP12-domains fused to GFP expressing HelaS3 cells. For each cell, DAPI granularity and GFP expression were determined as described in Materials and Methods on over than 50 cells. Blue and green filled circles correspond to individual non transfected and transfected cells, respectively. The linear regression curve is indicated in black. **E-** Representative images of chromatin condensates induced by TRIP12-IDR overexpression in HCT-116, U2OS and hTert-RPE1 cell lines by immunofluorescence. Nuclei were counterstained with DAPI. **F-** Comparison of isoelectric point (pI) and capacity to form chromatin condensates (slope) of the different TRIP12-GFP constructs. The pI of the different TRIP12 fragments was determined using ProtParam on Expasy website. The capacity to form chromatin condensates is indicated by the slope values obtained in Fig. 2D, 2F and 3C. **G-** Representative images of DNA organization in IDR-GFP deletion constructs in high expressing HelaS3 cells (left). The cytoplasmic leaking of 208-324-GFP and 325-445-GFP constructs is explained by the loss of NLS sequences. Determination of DNA granularity in function of GFP expression level for the different IDR-GFP deletion constructs (right). The DAPI granularity and GFP expression were determined as described in Materials and Methods on over than 50 cells. Blue and green filled circles correspond to individual non transfected and transfected cells, respectively. The linear regression curve is indicated in black. **H-** Comparison of isoelectric point (pI), capacity to form chromatin condensates (slope) and the length (in aa) of the different IDR-GFP deletion constructs. The pI and the length of the different TRIP12 fragments were determined using ProtParam on Expasy website. The capacity to form chromatin condensates was determined by slope values obtained in Fig. 3G. **I-** Graphical representation of MED1-IDR fused to GFP protein. The dark blue rectangle locates an artificial nuclear localization signals (NLS). Representative images of MED1-IDR-GFP expressing HelaS3 cells obtained by immunofluorescence. Nuclei were counterstained with DAPI. Cells with three different GFP intensities are represented. For each cell, DAPI granularity and GFP expression were determined as described in Materials and Methods on over than 50 cells. Blue and green filled circles correspond to individual non transfected and transfected cells, respectively. The linear regression curve is indicated in black.

We next calculated the isoelectric points (pI) of our different TRIP12 fusion proteins, which serves as an indirect indicator of its electric charge at physiological pH (∼7.4). Protein with pI > 7.4 can be expected to be positively charged in the cell, while proteins with pI < 7.4 will be negatively charged. The IDR of TRIP12 was the most positively charged domain (pI=10.76), and all the constructs that contain the IDR displayed a pI greater than 8.76 (Fig. 3F and Suppl. Fig. 2A). Indeed, many IDRs are positively charged due to the biophysical properties of their amino-acids^16^. The IDR of TRIP12 is enriched in serines (S)/lysines (K) and impoverished in leucines (L) (Suppl. Fig. 2A). Positive charges form patches along the IDR sequence of TRIP12 (Suppl. Fig. 2B). Interestingly, we determined a strong correlation between the charge of TRIP12 fragments and their capacity to induce the formation of chromatin condensates as indicated by their respective dose-response slope (Fig. 3F). This observation suggested that the formation of chromatin aggregates mediated by TRIP12 is likely driven by electrostatic forces.

Compared to IDRs of other proteins, the IDR of TRIP12 is a relatively large (440 aa) and covers ∼25% of the protein (Fig. 2B). We wondered whether the length of its IDR was an important parameter in the formation of chromatin condensates induced by TRIP12. To this aim, we generated a series of IDR deletion constructs and measured their ability to form chromatin condensates (Fig. 3G). Our analysis revealed that no specific sequences within the TRIP12-IDR was necessary for the formation of chromatin condensates. In contrast, we noted that the length of the IDR fragments was tightly correlated with their capacity to give rise to chromatin condensates, whereas the constructs displayed a similar pI (Fig. 3G and 3H). Indeed, IDR fragments of ∼100 aa lost their capacity to cause chromatin condensates. Beside, we noticed an enrichment in the nucleoli of most of IDR-GFP deletion constructs which can be explained by an enrichment of predicted nucleolar localization signals (NoLS) within the IDR (Suppl. Fig. 2C). For comparison, we measured the consequences of Mediator 1 (MED1)-IDR expression on chromatin organization. The biophysical features of the MED1-IDR (serine-rich, length of 627 aa and expected positive charge due to a pI = 10.01) (Suppl. Fig. 2D) were comparable to TRIP12-IDR (Suppl. Fig. 2B). Surprisingly, an over-expression of MED1-IDR-GFP at a comparable level to the one of TRIP12-IDR-GFP did not induce the formation of chromatin condensates (Fig. 3I).

Altogether, our results indicated that concentration, ionic charge and length of the TRIP12-IDR are critical parameters to drive the formation of chromatin condensates. Nevertheless, other unknown characteristics that seem unique to TRIP12 are likely required.

### The chromatin condensates induced by TRIP12-IDR are enriched in heterochromatin marks

We observed in Fig. 2 that an overexpression of TRIP12 induces a thickening of perinucleolar heterochromatin and the formation of large chromatin condensates. We therefore explored the content of these condensates by immunofluorescence using specific marks of constitutive (HP1α and H3K9me3) and facultative (EZH2, H3K27me3 and H2AK119ub) heterochromatin (Fig. 4A). We observed a strong colocalization of TRIP12-IDR-GFP staining with all these marks, demonstrating that TRIP12-induced chromatin condensates are mainly composed of heterochromatin. In contrast, two specific marks of transcriptionally-active euchromatin, the RNA polymerase II Serine 2- and Serine 5-phosphorylated forms were excluded from the condensates and rather localized to their periphery (Fig. 4B). Therefore, our results demonstrated that TRIP12-IDR overexpression leads to the formation of heterochromatin and consequently augments the volume occupied by heterochromatin regions.

**Figure 4:**
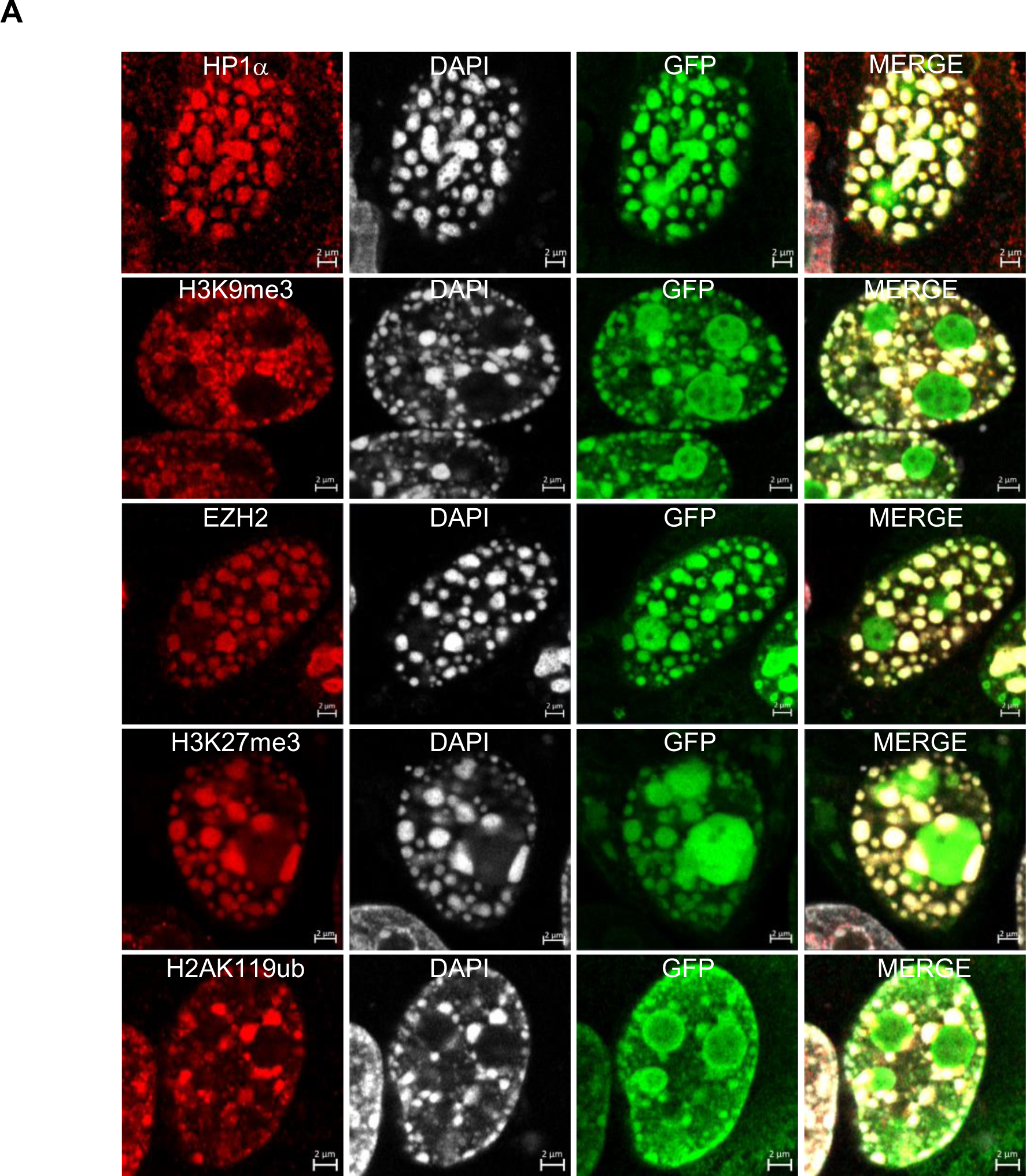
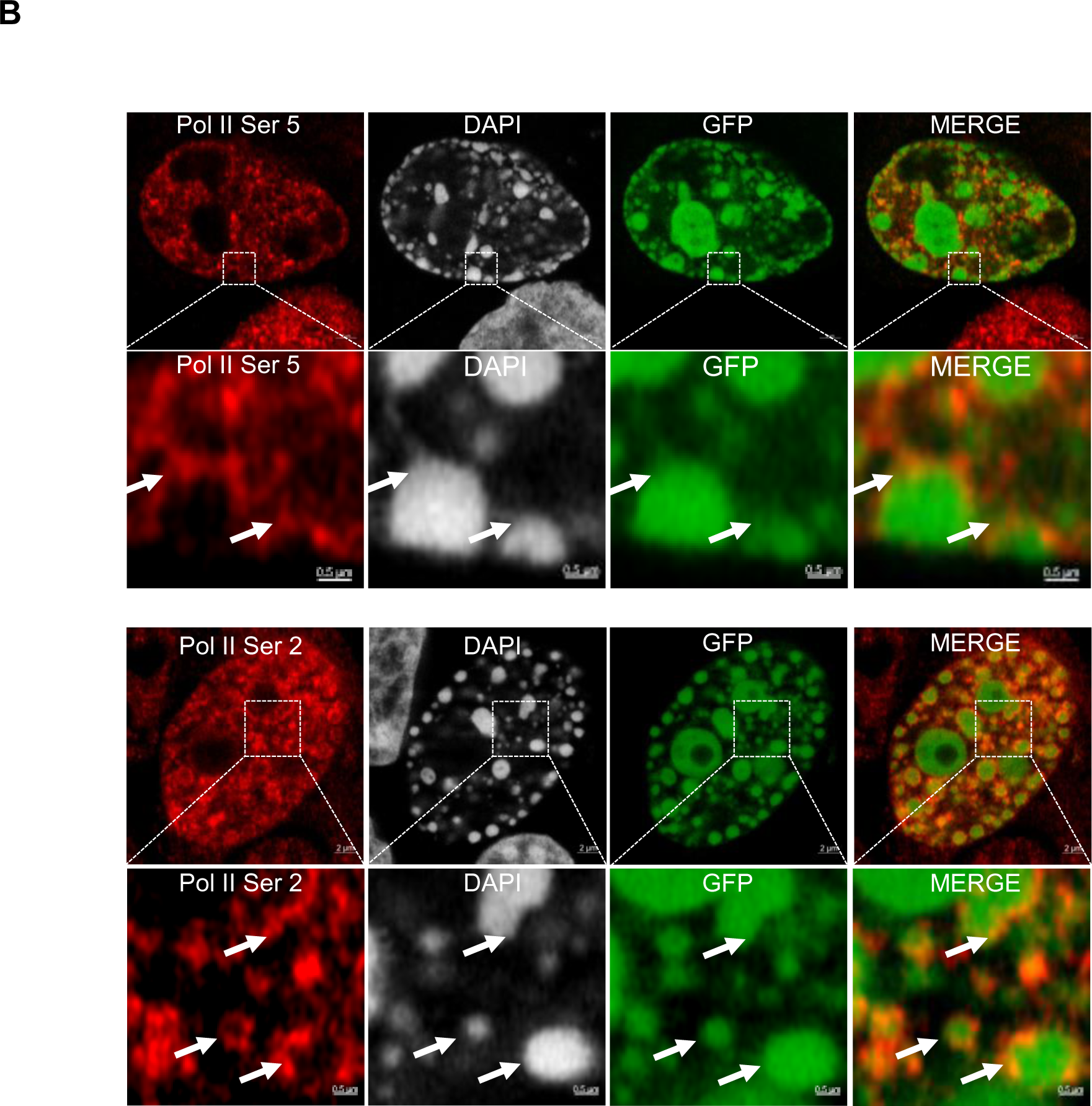
The chromatin condensates induced by TRIP12-IDR are enriched in heterochromatin marks. **A-** Representative images of HelaS3 cells transfected with IDR-GFP construct and localization of TRIP12-IDR mediated condensates and HP1α, H3K9me3, EZH2, H3K27me3 and H2AK119Ub heterochromatin marks by immunofluorescence. Nuclei were counterstained with DAPI. **B-** Representative images of localization of TRIP12-IDR mediated condensates and phosphorylated RNA polymerase II on serine 2 and serine 5. White arrows show the localization of RNA polymerase II staining at the periphery of the chromatin condensates. Nuclei were counterstained with DAPI.

### The formation of TRIP12-IDR mediated-chromatin condensates is a dynamic and reversible mechanism driven by polymer-polymer phase separation

We demonstrated that the formation of chromatin condensates is highly dependent of the expression level of TRIP12-IDR (Fig. 2). In cells with the highest level, TRIP12-IDR expression led to an ultimate level of compaction that allows the visualization of individual chromosomes with a characteristic centromeric CREST immunostaining suggesting a role of TRIP12 in mitotic chromosome condensation (Fig. 5A, top panel). Nevertheless, this profound reorganization of the genome did not modify the integrity of the nuclear membrane (Fig. 5A, bottom panel). We next set out to study the dynamics of chromatin condensate formation via an observation of individual TRIP12-IDR-GFP-expressing living cells by microscopy (Fig. 5B). We confirmed a direct correlation between the level of TRIP12-IDR expression and the formation of chromatin condensates within individual living cells (Fig. 5C). We further tested whether the chromatin condensates were reversibly or permanently formed upon TRIP12-IDR expression. To this aim, we generated an inducible VHL-NbGFP4-FLAG HelaS3 stable cell line expressing an intracellular anti-GFP nanobody (NbGFP4) coupled to a von Hippel Lindau (VHL)-E3 ubiquitin ligase to form a GFP degrader. First, we verified the efficient degradation of the TRIP12-IDR-GFP protein by the degrader (Fig. 5D). Second, we induced the formation of chromatin condensates by IDR-GFP transfection followed by the expression of the GFP-degrader. After 24h, we measured a significant decrease of GFP intensity correlated to a diminution of chromatin condensates, demonstrating the reversibility of chromatin condensates formation triggered by TRIP12 expression (Fig. 5E and 5F). In parallel, we explored the kinetics by monitoring individual living cells with chromatin condensates for 13h after the expression of the GFP-degrader by microscopy. We observed that an IDR-GFP degradation diminishes the presence of chromatin condensates within a few hours (Fig. 5G and 5H), confirming the reversibility of chromatin condensates formation induced by TRIP12.

**Figure 5:**
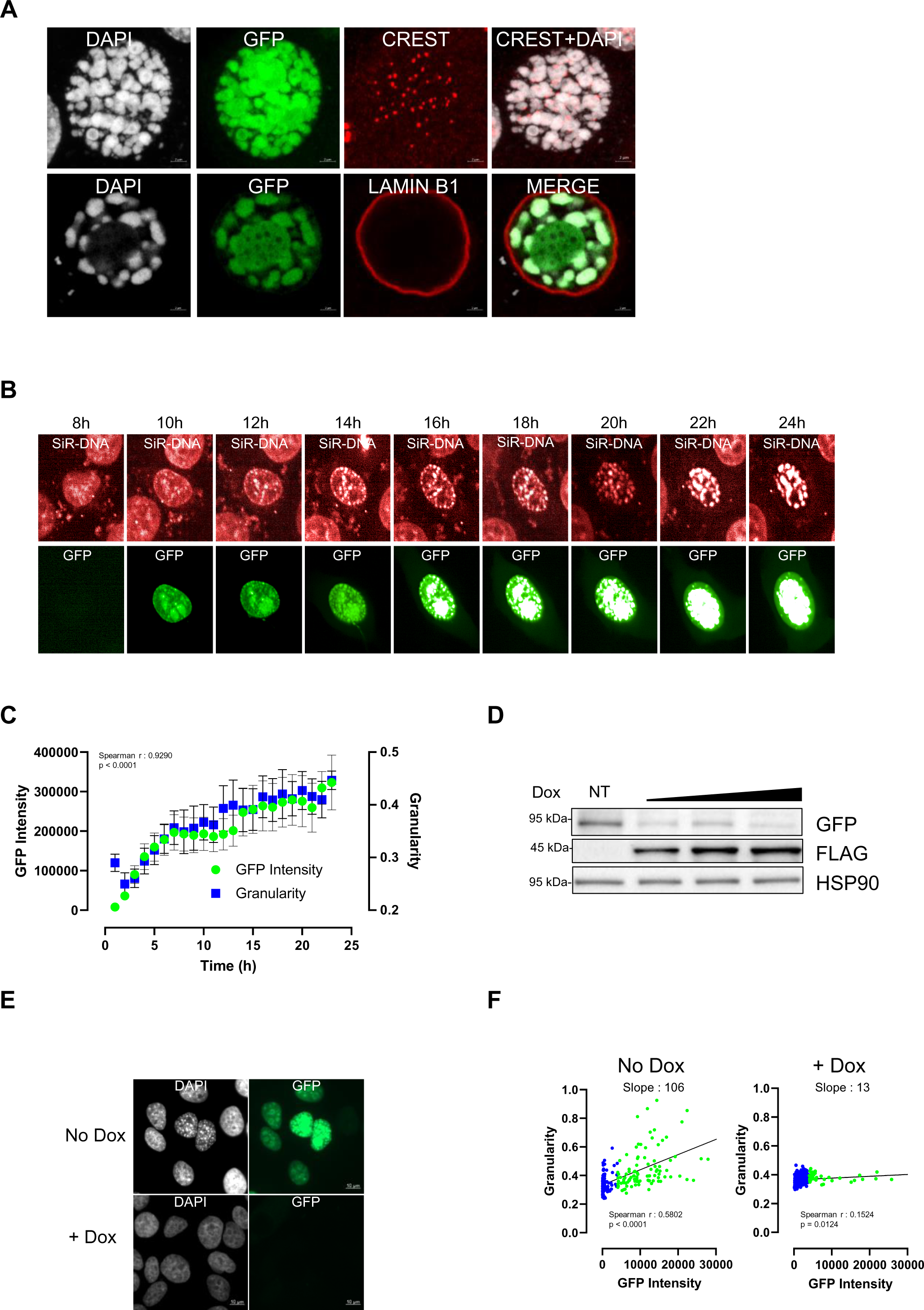
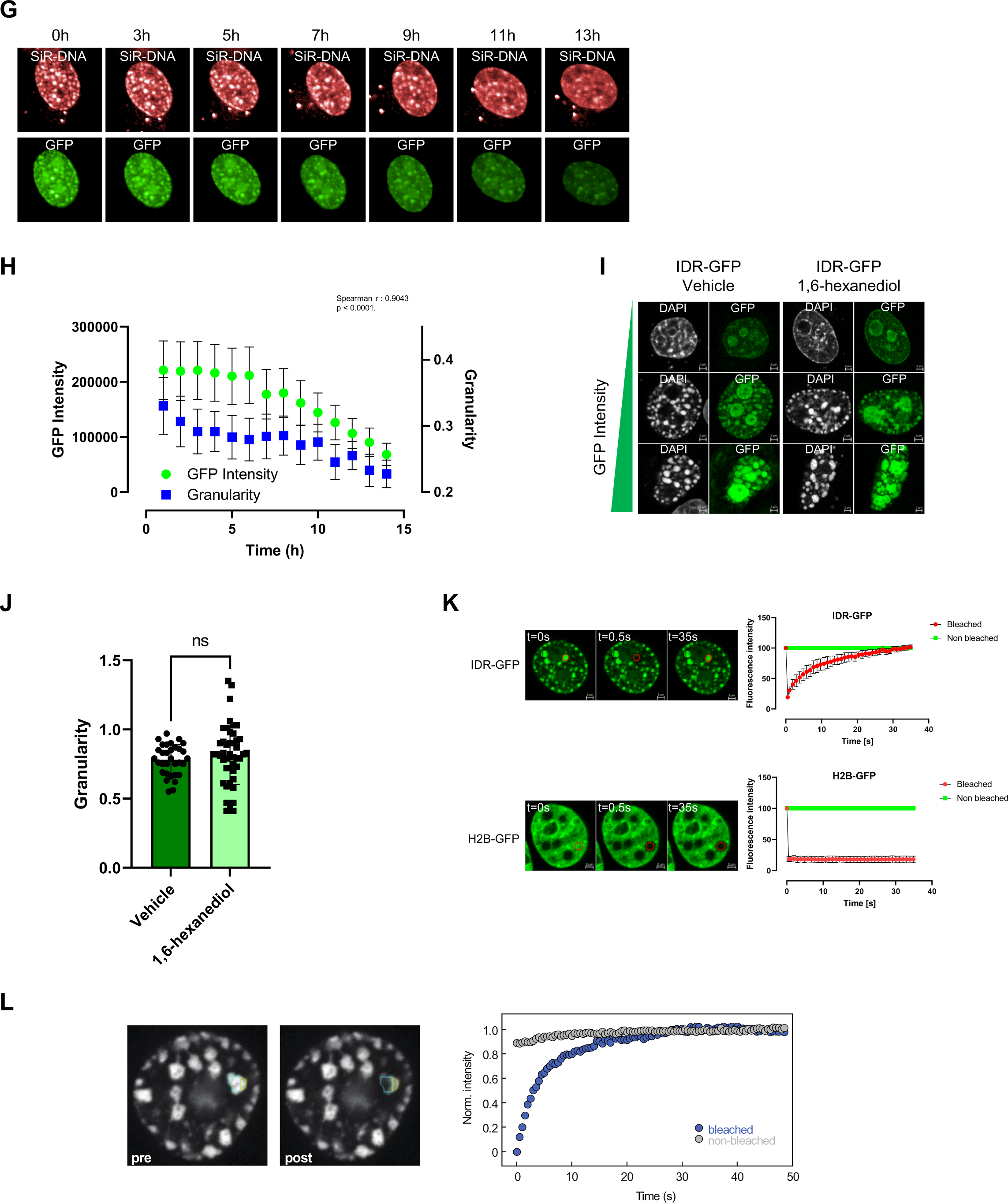
The formation of TRIP12 induced-chromatin condensates is a dynamic and reversible mechanism driven by polymer-polymer phase separation. **A-** Visualization of CREST (top) and LAMIN B1 (bottom) and chromatin condensates in TRIP12-IDR expressing HelaS3 cells by immunofluorescence. Nuclei were counterstained with DAPI. **B-** Representative images of chromatin condensate formation and GFP expression in IDR-GFP expressing HelaS3 cells by live cell microscopy. Images were acquired every hour for 24 h after transfection. For the clarity of the figure, representative image of a cell 8 h after transfection and every two hours are represented. Nuclei were counterstained with SiR-DNA-647 nm. **C-** Quantification of GFP expression and DNA granularity in IDR-GFP expressing HelaS3 cells obtained by live cell microscopy. Images were acquired every hour for 24 hours by live cell confocal microscopy. GFP intensity and DNA granularity were determined using Columbus and FIJI software. Results are expressed as mean ± SEM of at least five independent cells. **D-** Expression of IDR-GFP, FLAG-tagged anti-GFP degrader in VHL-NbGFP4-FLAG expressing HelaS3 cells determined by Western blot 24 h after increasing doses of doxycycline (0.25, 0.5 and 1 µg/ml). The immunoblots are representative of three independent experiments. HSP90 protein level was used as loading control. **E-** Representative images of VHL-NbGFP4-FLAG expressing in HelaS3 cells transfected with IDR-GFP construct in the presence or not of doxycycline (1 µg/ml) for 24 h obtained by immunofluorescence. Nuclei were counterstained with DAPI. **F-** Determination of DNA granularity in function of IDR-GFP expression level in VHL-NbGFP4-FLAG expressing in Hela cells transfected with IDR-GFP construct in the presence or not of doxycycline (1 µg/ml) for 24 h or vehicle. For each cell, DAPI granularity and GFP expression were determined as described in Materials and Methods on over than 50 cells. Blue and green filled circles correspond to individual non transfected and transfected cells, respectively. The linear regression curve is indicated in black. **G-** Representative images of DNA condensates and IDR-GFP expression in VHL-NbGFP4-FLAG expressing HelaS3 cells transfected with IDR-GFP construct every hour (for 13 h) after addition of doxycyclin (1 µg/ml) by cell live microscopy. DNA is stained with SiR-DNA-647 nm. **H-** Quantification of DNA granularity and GFP intensity in VHL-NbGFP4-FLAG expressing HelaS3 cells transfected with IDR-GFP construct every hour (for 13 h) after addition of doxycyclin (1 µg/ml) by cell live microscopy. ± SEM of at least five independent cells. **I-** Representative images of DAPI organization in IDR-GFP expressing HelaS3 cells in the presence of 1,6-hexanediol 0,5% or vehicle for 18 h. Cells with three different GFP intensities are represented. Nuclei were counterstained by DAPI. **J-**Determination of DAPI granularity in IDR-GFP high expressing HelaS3 cells in the presence of 1,6-hexanediol 0,5% or vehicle for 18 h as described in Materials and Methods section. **K-** IDR-GFP mobility in DNA condensates by FRAP analysis. H2B-dsRed expressing HelaS3 cells were transfected with IDR-GFP construct or H2B-GFP used as control. After 24 h, GFP fluorescence was photo-bleached using a FRAP module of confocal microscope. The recovery of fluorescence was measured every second for 35 sec. Representative images of IDR-GFP and H2B-GFP expressing cells with bleached and non-bleached areas at indicated times are represented (left). The graph represents the mean ± SD of GFP fluorescence intensity obtained from four independent cells. **L-** IDR-GFP mobility in DNA condensates by half-bleached FRAP analysis. HelaS3 cells were transfected with IDR-GFP construct. After 24 h, GFP fluorescence in a half of a condensate was photo-bleached using a FRAP module of confocal microscope. The recovery of GFP fluorescence was measured every second for 50 sec. Representative images of an IDR-GFP expressing cell with bleached and non-bleached areas before and just after bleaching are represented (left). The graph represents the fluorescence intensity in a bleached and a none bleached condensate in one cell.

Chromatin condensates induced by IDR-containing proteins can be generated by two main non-exclusive highly dynamic processes: liquid-liquid phase separation (LLPS) or polymer-polymer phase separation (PPPS)^19^. Both processes can be driven by electrostatic or hydrophobic interactions. To determine the contribution of the latter, cells were treated with 1,6-hexanediol (1,6-HD), an aliphatic alcohol that impairs hydrophobic interactions. This treatment neither impacted the formation of chromatin condensates (Fig. 5I) nor the DAPI granularity (Fig. 5J). Moreover, we analyzed the dynamics of TRIP12-IDR-GFP in chromatin condensates by FRAP. Compared to H2B-GFP, which showed no visible recovery, TRIP12-IDR-GFP fully recovered within tens of seconds, indicating that TRIP12-IDR-GFP interacts transiently with chromatin (Fig. 5K). To test if condensates are surrounded by an interface that restricts molecular transport, which would be indicative of LLPS, we performed half-FRAP experiments in which we photo-bleached one half of the condensates and measured the fluorescence intensity in both halves^30,37^ (Fig. 5L). The intensity in the non-bleached half did not show a pronounced decrease, indicating that the recovery in the bleached half originated mostly from exchange of molecules between the condensate and the surrounding nucleoplasm and only to a small extent from internal mixing between both halves of the condensate. Accordingly, these experiments indicated that the half-bleached condensates do not contain an interfacial barrier.

Together with the results shown in Fig. 3, we conclude that chromatin condensates induced by TRIP12-IDR are highly dynamic and are likely formed by PPPS involving electrostatic forces.

### Consequences of TRIP12-IDR mediated-chromatin condensates on cellular growth, chromatin accessibility and genome expression

Next, we investigated whether IDR-mediated-chromatin condensates could influence cellular growth. IDR-GFP and H2B-GFP transfected HelaS3 cells were monitored for 30 h by live cell microscopy and classified into three categories depending on their level of GFP expression. Interestingly, during the observation period, low and moderate IDR-GFP expressing cells underwent only one cell division, in contrast to H2B-GFP expressing cells that divided twice for most of them. Moreover, the vast majority of cells expressing a high level of IDR-GFP died during the first 20 h after transfection (Fig. 6A). A subsequent analysis of the cell cycle distribution showed that all IDR-GFP cell categories accumulate in late S and G_2_ phases comparatively to H2B-GFP cells (Fig. 6B). These results indicated that the presence of chromatin condensates caused by the TRIP12-IDR perturbs the cell cycle progression via a late S/G_2_ arrest. To test whether DNA replication defects could explain the late S/G_2_ cell enrichment, IDR-GFP expressing cells were cultured in the presence of EdU. As seen in Fig. 6C, the incorporation of EdU in the condensates suggested that the chromatin compaction caused by TRIP12 did not drastically alter DNA replication and that the late S/G_2_ arrest was likely due to other functional alterations.

**Figure 6:**
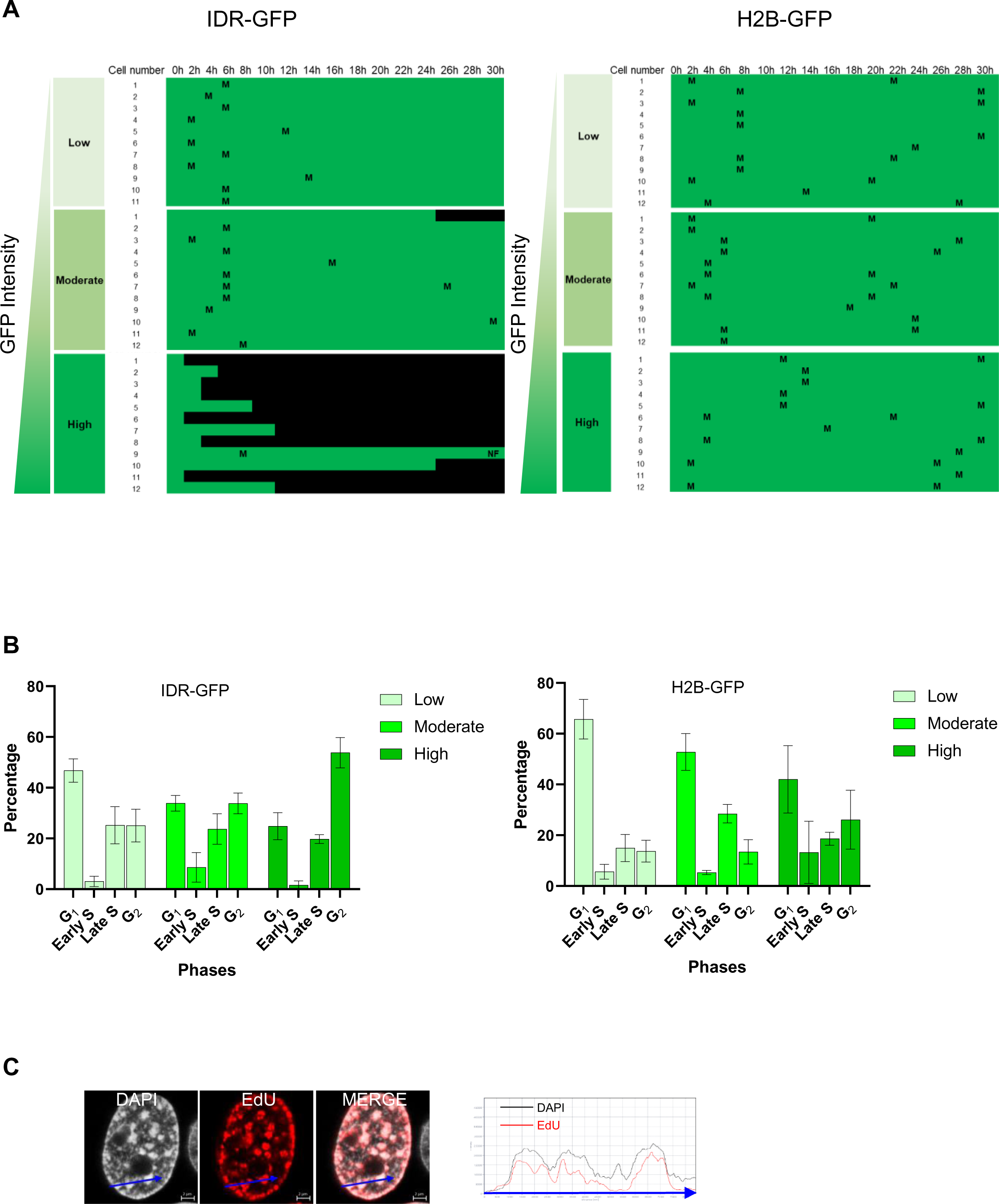
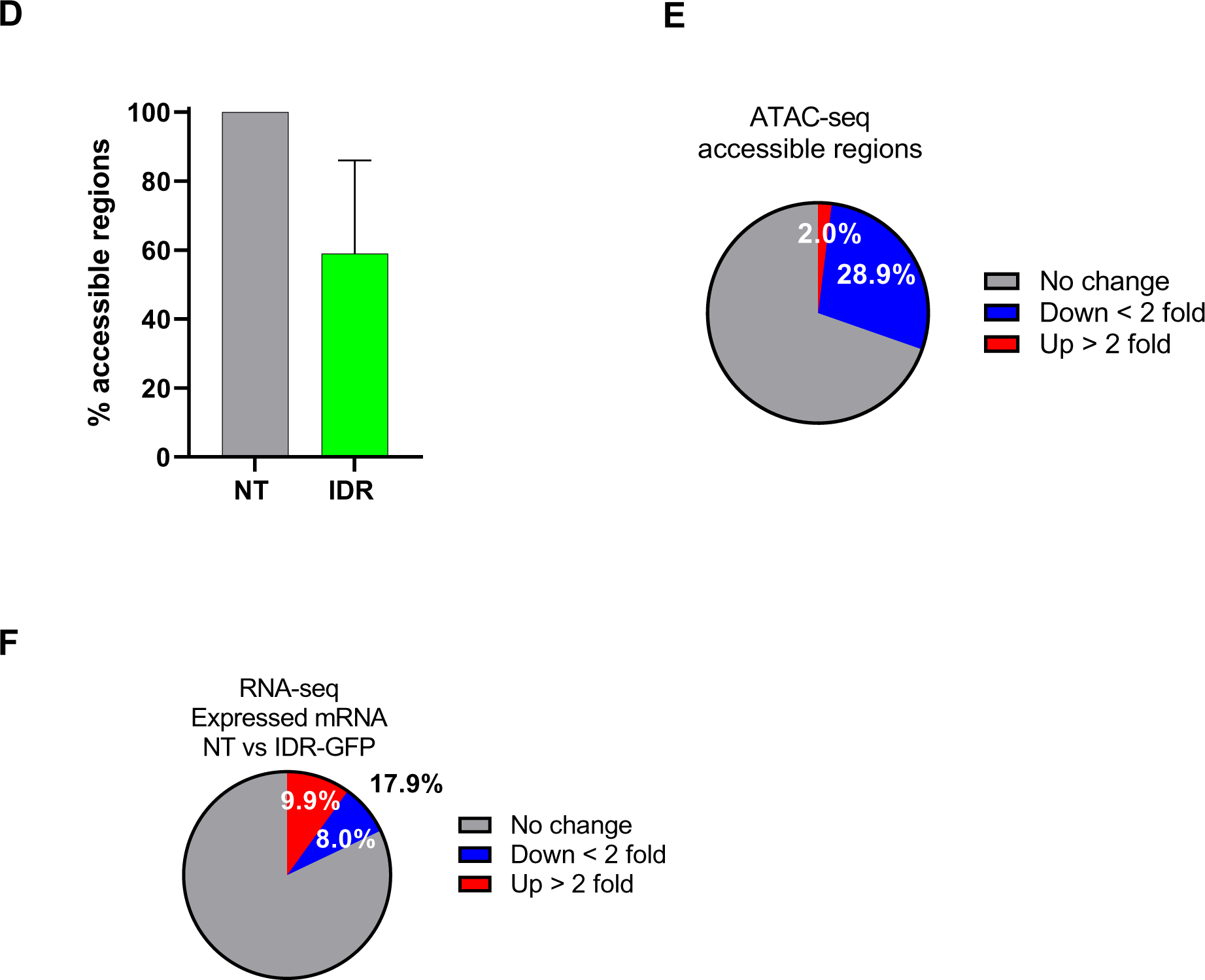
Consequences of TRIP12-IDR mediated-chromatin condensates on cellular growth, chromatin accessibility and genome expression. **A-** Cellular fate of IDR-GFP (left) and H2B-GFP (right) transfected HelaS3 cells. Twenty-four hours after transfection, fluorescence of IDR-GFP transfected cells was determined using Incucyte apparatus to classify cells in three categories: low, moderate and high. Eleven to twelve cells per category were monitored for 30 h every 2 h. Mitotic divisions were annotated with a M and nuclear fusion with a NF. Green bars represent living cells whereas black bars represent dead cells. **B-** Distribution in the cell cycle of IDR-GFP (left) and H2B-GFP (right) transfected HelaS3 cells. Twenty-four hours after transfection, the position in the cell cycle was determined by immunofluorescence using CYCLIN A and EdU staining as described in Materials and Methods. The graph represents the percentage of cells in the different phases of the cell cycle expressed as mean (± SEM) of at least three independent experiments. **C-** Representative image of EdU incorporation in IDR-GFP-mediated chromatin condensates (left). HelaS3 cells were transfected with IDR-GFP construct. Twenty-four hours after transfection, cells were incubated with EdU (1 µM) for 20 min. Incorporation of EdU was visualized by Click-It™ followed by confocal microscopy. Nuclei were counterstained with DAPI. The graph represents the intensity of fluorescence along the arrow (right). **D-** Determination of genomic accessible regions in IDR-GFP expressing cells by ATAC-seq experiments. Only merged regions with a Max tags > 50 were considered. Results were obtained from two independent experiments and expressed as a percentage ±SEM of genomic accessible regions compared to non-transfected cells (set as 100%). **E-** Percentage of accessible regions in IDR-GFP expressing HelaS3 cells with a lower (Down, fold change<2), higher (Up, fold change>2) and unchanged accessibility compared to non-transfected cells (set as 100%). Results were obtained from two independent experiments. **F-** Percentage of down (fold change<2) and up-regulated (fold change>2) mRNA in IDR-GFP expressing HelaS3 cells compared to non-transfected cells by RNA-seq approach. Results were obtained from two independent experiments.

The accessibility of the genome to transcriptional machinery is tightly controlled by chromatin compaction which can be estimated by ATAC-seq approaches. As expected, we found that the number of accessible regions was dramatically reduced in IDR-GFP high expressing cells (Fig. 6D). A more detailed analysis revealed a high percentage (28.9%) of regions for which the accessibility was decreased compared to a low percentage (2%) of regions for which the accessibility was increased (Fig. 6E). In parallel, a transcriptome analysis by RNA-seq on the cells that were used for ATAC-seq revealed that among 8906 detectable mRNAs, the expression of 17.9% (n=1 598) of them was statistically modified by an over-expression of IDR-GFP protein. More precisely, 8.0% (n=713) were downregulated and 9.9% up-regulated (n=885) more than two-fold (Fig. 6F). We could not establish a statistical correlation between the level of gene accessibility and their mRNA expression Nevertheless, these results clearly indicated that an overexpression of the IDR of TRIP12 greatly alters the accessibility of the genome as well as its expression.

### TRIP12 overexpression inhibits the accumulation of MDC1 protein on DNA double strand breaks via its IDR

It is well-documented that chromatin compaction is a critical element in the efficiency of DDR pathways^1,2^. Interestingly, we noticed that the protein Mediator of DNA damage checkpoint 1 (MDC1), a key protein in the detection of DNA double strand breaks (DSB)^38^, is among the most enriched proteins in the proxisome of TRIP12 (Fig. 1C and Suppl. Table 3). Indeed, in response to DSB, MDC1 forms foci by recognizing the phosphorylated serine S139 of histone H2A.X (γH2A.X). In turn, MDC1 participates in the recruitment of crucial proteins of the DNA damage response (DDR) pathways such as P53 Binding Protein (53BP1) (NHEJ) and Breast Cancer 1 (BRCA1) (HR)^38^. An overexpression of NBirA*-FLAG-TRIP12 in X-ray-irradiated cells dramatically inhibited the formation of MDC1 foci in all cell cycle phases unlike an overexpression of NBirA*-FLAG-EGFP-NLS used as control (Fig. 7A and 7B). Of note, the overexpression of NBirA*-FLAG-TRIP12 did not alter the number of γH2AX foci, meaning that TRIP12 does not affect the early detection of DNA breaks upstream of MDC1 accumulation (Suppl. Fig. 3A and 3B). To identify the domain(s) responsible for MDC1 accumulation, HelaS3 were transfected with different GFP-constructs and subsequently irradiated. Similarly, an overexpression of TRIP12-GFP drastically impeded MDC1 foci formation (Fig. 7C and 7D). This inhibition was also measured with TRIP12 catalytic mutants (TRIP12-GFP C1959A and ΔHECT-GFP) but not with a construct lacking the IDR (ΔIDR-GFP), suggesting that this domain is required. Indeed, the overexpression of IDR-GFP alone was sufficient to remarkably inhibit the formation of MDC1 foci in a dose-dependent manner (Fig. 7E and 7F) even in IDR-GFP low expressing cells in which chromatin modifications are subtle (Fig. 2C). Moreover, the inhibition of MDC1 accumulation by TRIP12 via its IDR was kinetically confirmed after UV-laser micro-irradiation (Fig. 7G and 7H). Indeed, a prior expression of full-length TRIP12 fused to the RFP reporter protein dramatically reduced the recruitment of GFP-MDC1 to the irradiated zone when an expression of RFP-IDR alone totally abolished it.

**Figure 7:**
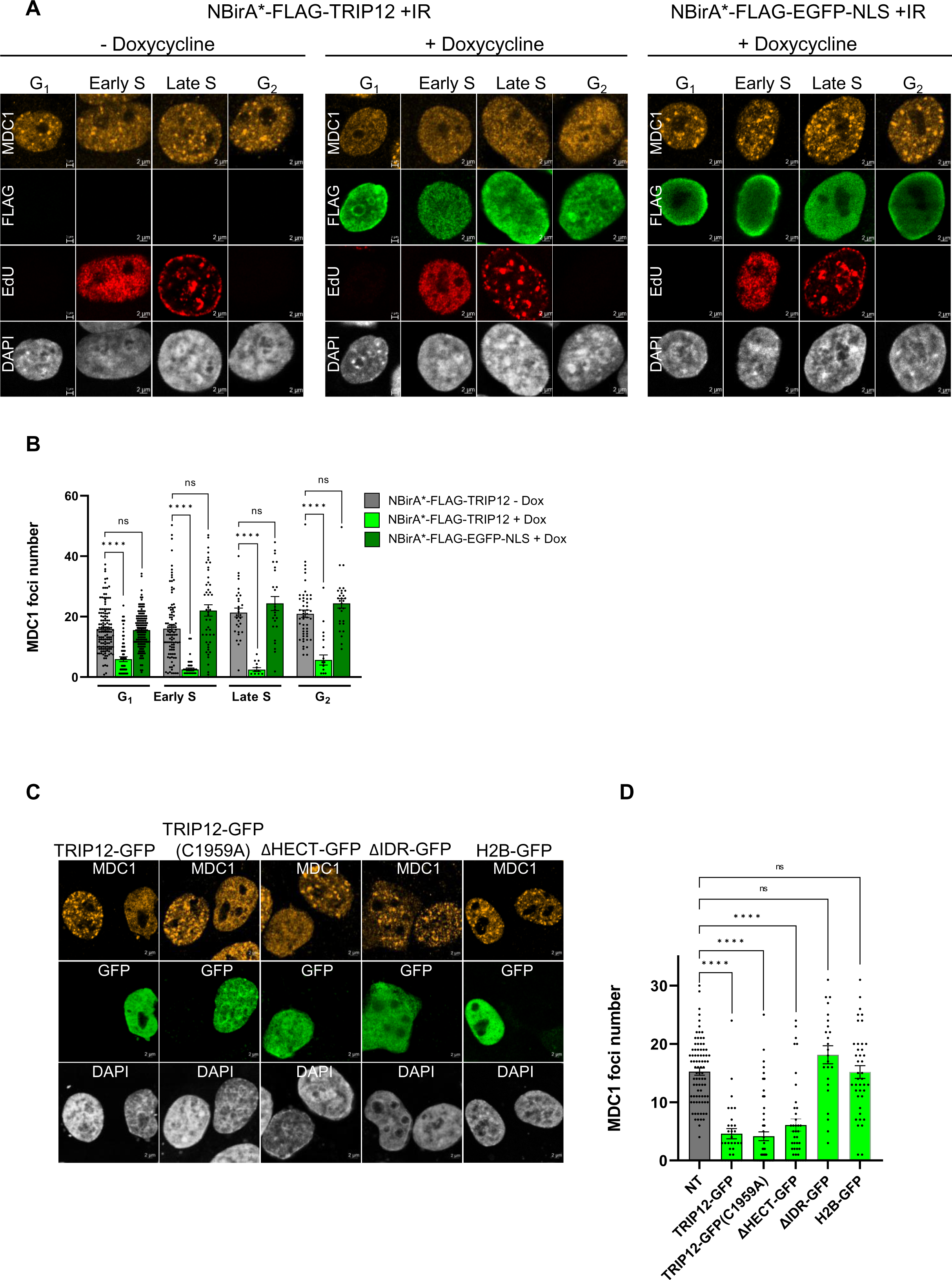
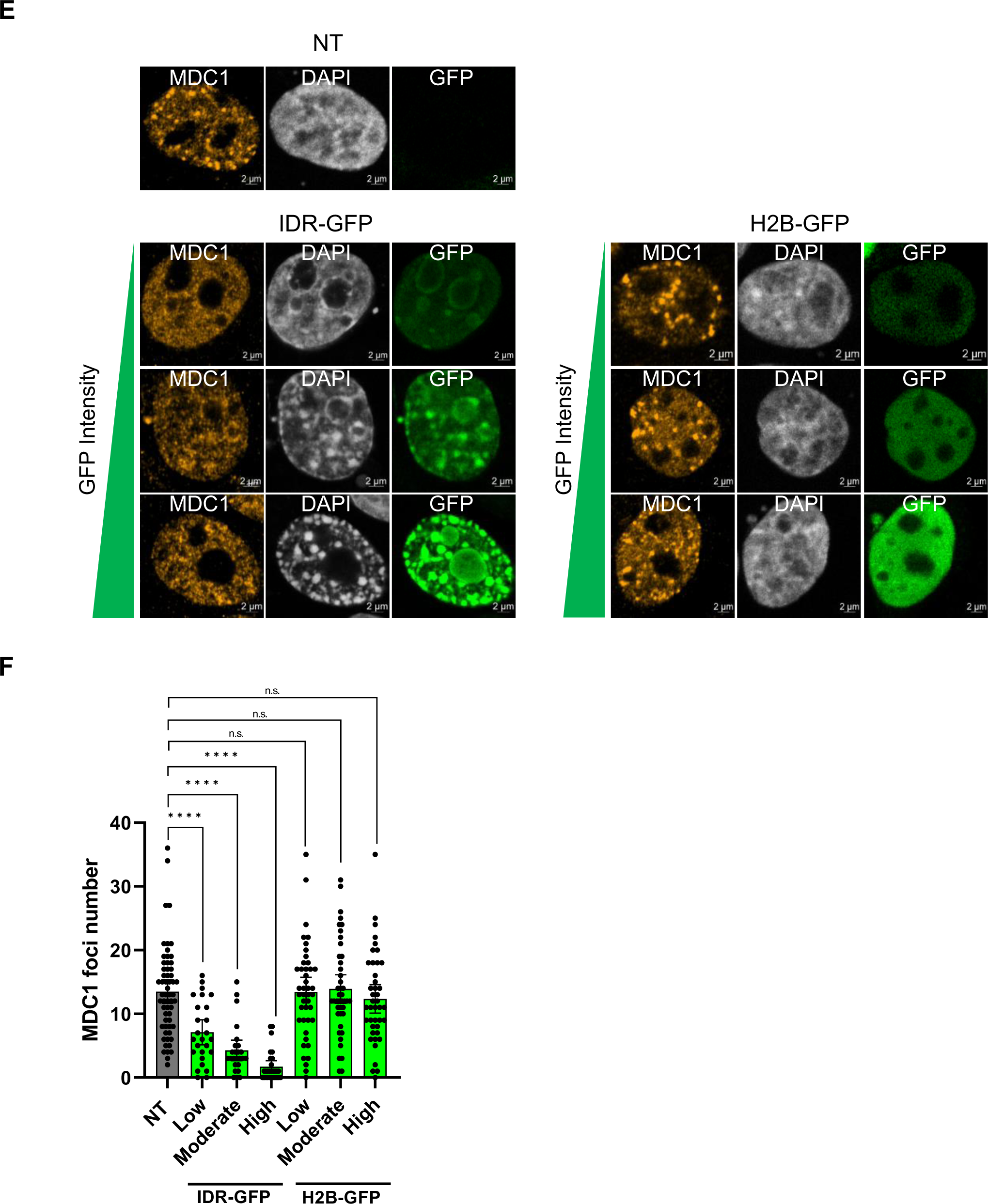
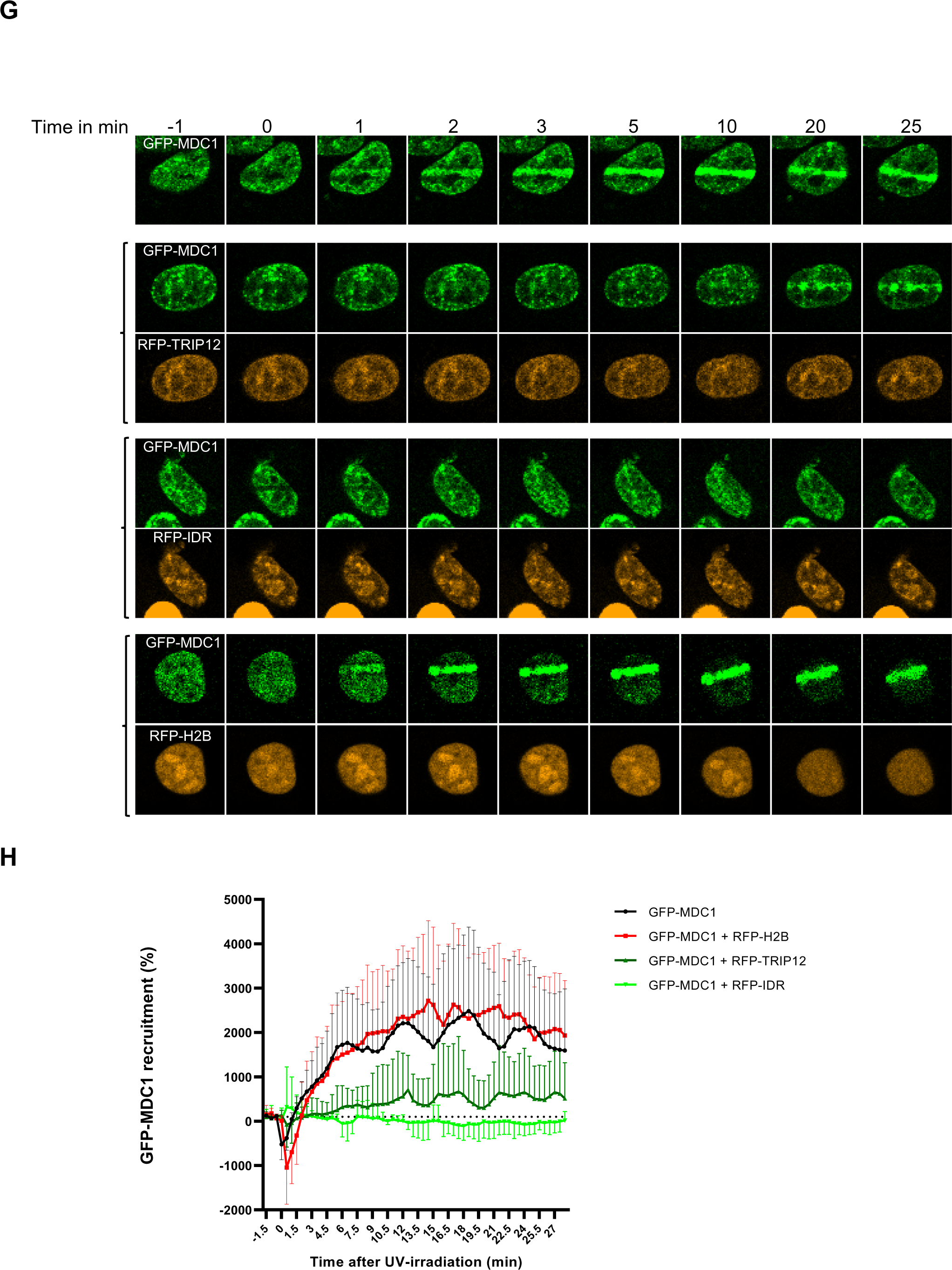
TRIP12 overexpression inhibits the accumulation of MDC1 protein on DNA double strand breaks via its IDR. **A-** Representative images of MDC1 foci, EdU and FLAG in irradiated NBirA*-FLAG-TRIP12 and NBirA*-FLAG-EGFP-NLS cell lines treated or not with doxycycline (1 µg/ml for 48 h) during cell cycle phases by immunofluorescence. Cells were X ray-irradiated (1 Gy) and treated with EdU 1 h and 20 min, respectively, before the end of 48 h-doxycycline treatment. The determination of the cell cycle phase is described in Materials and Methods. Nuclei were counterstained with DAPI. **B-** Quantification of MDC1 foci in irradiated NBirA*-FLAG-TRIP12 and NBirA*-FLAG-EGFP-NLS cells treated or not with doxycycline during the different phases of the cell cycle. Each dot corresponds to one cell. Results are expressed as mean ±SEM obtained from independent experiments determined by confocal microscopy and FIJI macro command. **C-** Representative images of MDC1 foci and GFP staining in irradiated HelaS3 cells transfected or not indicated GFP constructs by immunofluorescence. Nuclei were counterstained with DAPI. **D-** Quantification of MDC1 foci in irradiated HelaS3 cells transfected or not indicated GFP constructs. Each dot corresponds to one cell. Results are expressed as mean ± SEM obtained from independent experiments determined by confocal microscopy and FIJI macro command. **E-** Representative images of MDC1 foci and GFP by immunofluorescence in X ray-irradiated HelaS3 cells expressing IDR-GFP or H2B-GFP. Twenty-three hours after transfection, cells were X-ray irradiated (1 Gy). A non-transfected cell and cells with three different GFP intensities are represented. Nuclei were counterstained with DAPI. **F-** Quantification of MDC1 foci in irradiated IDR-GFP and H2B-GFP expressing HelaS3 cells depending on GFP expression. IDR-GFP expressing cells were classified in low, moderate and high depending on the intensity of GFP expression. Each dot corresponds to one cell. Results are expressed as mean ± SEM obtained from independent experiments determined by confocal microscopy and FIJI macro command. **G-** Representative images of GFP and RFP fluorescence in HelaS3 cells expressing the indicated constructs and UV-laser (405 nm) micro-irradiated (time 0) at the indicated times in min by live cell microscopy. For clarity, not all the time points performed in the experiments are represented on the figure. **H-** Quantification of GFP-MDC1 recruitment kinetics in UV-laser micro-irradiated HelaS3 cells transfected with the indicated constructs at the indicated times. The results are expressed as mean ±SD obtained from 10 cells of two independent experiments. The dotted line represents 100% of recruitment.

These results clearly demonstrated that an overexpression of TRIP12 affects DNA damage response through the action of its IDR by inhibiting the accumulation of the DDR-key protein MDC1 on DNA breaks.

## DISCUSSION

Up to now, the roles attributed to the E3 ubiquitin ligase TRIP12 have exclusively implicated its C-terminal-ubiquitin ligase activity. Herein, we biophysically characterized for the first time a functional role for the TRIP12 intrinsically disordered N-terminal domain on chromatin compaction with major consequences on several important biological processes such as DDR.

To begin, we established the first interactome of TRIP12 used as a bait by BioID. It confirmed the pleotropic implication of TRIP12 in chromatin compaction and organization networks. Numbers of partners were found in common with interactomes in which TRIP12 was used as a prey (i.e.: BRD4^39^) which enhanced our confidence in our identified proteins. One advantage of this approach is its capacity to identify (without discriminating them) not only protein interacting partners but also ubiquitinated substrates, for which the interaction is extremely labile^14,40^. Therefore, our analysis provides the scientific community with an exhaustive list of candidate proteins that will help, with further investigations, to better understand the complex mechanisms of TRIP12-mediated chromatin regulation.

We next demonstrated that an overexpression of TRIP12 induces the formation of chromatin condensates. More precisely, we identified the IDR as responsible domain for this compaction. We further showed that the nuclear concentration of TRIP12 or its IDR is a critical parameter in the degree of chromatin compaction. Also, we discovered that the electric charge of the IDR (pI=10.76) is likely important for the capacity of TRIP12 to form chromatin condensates. These findings indicate that chromatin condensation induced by TRIP12 is driven by electrostatic forces. The IDR of TRIP12 is enriched in serine, the third most frequent residue found in disordered regions^41^. Positive charges are heterogeneously spread on the TRIP12-IDR sequence and form patches. It is known that the distribution of positive and negative charges within IDRs is important to exert their function^42^. Electric charge is a major parameter in chromatin condensate formation by TRIP12 but not the only one. Indeed, the length of the IDR is also an important parameter as it loses its capacity to form condensates when shrunk to ∼100 aa likely by reducing the number of electric charges. Chromatin condensates can be driven by the two non-exclusive biophysical processes PPPS and LLPS^19^. Among them, LLPS has been more often implicated in heterochromatin and the regulation of the transcriptional machinery, implicating IDR-proteins such as BRD4 or MED1^17^, respectively. PPPS and LLPS are highly dependent on protein concentration, which must reach a saturating concentration to form condensates. We showed that chromatin condensates formed by the TRIP12-IDR are likely driven by PPPS involving electrostatic forces. Altogether, our study reveals that concentration, positive electric charge and length of the IDR of TRIP12 are essential parameters to lead to the formation of chromatin condensates. Although almost half of the proteome possesses IDRs, only 10-20% of these proteins contain a disordered region that covers more than 25% of the protein size as TRIP12-IDR does^16^. This feature combined with a length of 440 aa (50 kDa) makes the IDR of TRIP12 very unique. Notably, the MED1-IDR that displays similar length (600 aa) and electric charge (pI=10.01) does not induce chromatin condensates when expressed at the same level as TRIP12-IDR. This raises the question whether this property is specific to TRIP12. Similar macroscopic chromatin condensates driven by PPPS were observed after overexpression of the PHD finger protein in ovarian cancer (SPOC1) protein. SPOC1-mediated chromatin compaction involves two structured protein domains and also impacts DDR^43–45^.

The formation of heterochromatin is driven by a multi-step process that includes post-translational histone modifications by protein complexes and the subsequent recruitment of reader proteins. For instance, HP1α is known to induce by oligomerization the formation of heterochromatin in vitro^46^. Nevertheless, it is suggested that, in a cellular context, HP1α is not sufficient to form such a level of chromatin compaction which requires other proteins^47^. In our study, we discovered that TRIP12 expression induces the formation of chromatin condensates enriched in specific heterochromatin marks. It is therefore tempting to speculate that TRIP12 participates in the formation of heterochromatin via electrostatic interactions. The targeting of TRIP12 to specific genomic regions could be ensured by the PRC complexes. To support this idea, our BioID analysis revealed numbers of PRC complex proteins in the proxisome of TRIP12. Moreover, several reports identified TRIP12 in the interactome of PRC complexes proteins^11,12^.

The level of chromatin compaction has been involved in several important biological processes. A proper chromosome compaction is a crucial step for mitotic cell division. Herein, we observed that chromatin compaction induced by an expression of IDR-GFP perturbs the cell cycle and leads to an accumulation of cells in late S/G_2_ phase and a subsequent death during mitosis. A finely-tuned expression of TRIP12 seems required to complete mitotic division. Therefore, it is likely that TRIP12, via its IDR properties, participates in the proper condensation of chromosomes prior to mitosis. This is in accordance with our previous study in which we showed that TRIP12 is tightly controlled during the cell cycle and highly expressed in late S and G_2_ phases^15^. Moreover, we noticed that a depletion of TRIP12 results in alterations of chromosome segregation. DNA replication efficiency during S phase is tightly dependent on chromatin assembly^48^. Herein, chromatin condensates induced by TRIP12-IDR do not inhibit the incorporation of EdU, suggesting that they do not alter DNA replication. Nevertheless, to confirm this claim, the speed of DNA replication forks in the presence or absence of the TRIP12-IDR should be measured.

Gene transcription is another important molecular process that is tightly dependent on chromatin compaction level. Genome accessibility to the transcriptional machinery dictates what genes must be transcribed into mRNA. We measured the impact of TRIP12-IDR induced-condensates on genome accessibility and gene transcription by ATAC-seq coupled to RNA-seq. We noticed a dramatic decrease in the number of accessible genomic regions, coupled to profound alterations of the transcriptome, indicating a direct implication of TRIP12-IDR in transcriptional regulation. Deciphering the molecular mechanisms that govern this alteration is a remaining task. The exclusion of active RNA polymerase II at the periphery of chromatin condensates induced by TRIP12 that we observed in this study could be a straightforward explanation. The histone variant H2A.Z has been strongly correlated with transcription control^49^. Interestingly, this histone variant is the most enriched protein identified in our BioID analysis. Therefore, the mechanistic relationship between TRIP12 and H2A.Z could provide deeper insights into the implication of TRIP12-IDR in the control of transcription.

The efficiency of the DDR greatly depends on chromatin compaction. Nuclear F-actin and myosin drive the re-localization of heterochromatic breaks in relaxed chromatin^50^. Up to now, TRIP12 has been implicated in the inhibition of NHEJ and Alternative End joining (Alt-EJ) DDR pathways via the action of its HECT catalytic domain by inducing the ubiquitination of RNF168 and PARP1, respectively^4,5^. Here, we discovered that the IDR of TRIP12 inhibits the formation of MDC1 foci at DSBs in a dose-dependent manner. MDC1 is a central protein in the DDR as it participates in the recruitment and the accumulation of 53BP1 and BRCA1^51^. MDC1 is among the most enriched proteins in the TRIP12 BioID proxisome which is in accordance with a recent study that identified TRIP12 as a main interactant of MDC1^52^. Nevertheless, the molecular relationships that exist between MDC1 and TRIP12-IDR are still only partially known. We can speculate that chromatin compaction induced by TRIP12-IDR at DSBs creates a steric hindrance for MDC1 recruitment and accumulation. Another explanation would be that TRIP12-IDR physically interacts with MDC1. Consequently, an overexpression of TRIP12 and its fixation along the genome would titrate MDC1 and impedes its recruitment at DSBs. H2AX phosphorylation on Ser 139 (γH2AX) is associated with modifications of chromatin structure^53^. By modifying the structure of chromatin, TRIP12, via its IDR, could also trigger a genome-wide H2AX phosphorylation that would titrate the free-MDC1 pool onto the chromatin as MDC1 specifically recognizes H2AX (Ser 139) phosphorylation by its BRCA1 C-Terminus (BRCT) domain ^38^.

The Trip12 mRNA is overexpressed in numerous cancers and notably in pancreatic cancers in which we (Personal data) and others showed a heterogeneous overexpression^3,36^. Several studies have reported the implication of TRIP12 via its HECT catalytic activity in the sensitivity of cancer cells to chemotherapies^4,5^. Given the inhibitory effect on MDC1 foci formation, it is likely that the IDR of TRIP12 simultaneously participates in the chemosensitivity of cancer cells. Finally, numerous studies have reported an association between Trip12 mutations and intellectual deficiencies such as Clark-Baraister syndrome^21,54^. The molecular mechanisms involved are currently unknown. Numbers of non-sense Trip12 mutations identified in patients produce truncated proteins corresponding to the IDR alone^20^. Based on our discoveries, alterations of chromatin condensation could constitute the premise for a better understanding of those diseases.

## DATA AVAILABILITY

Data are deposited in a public repository as indicated in Materials and Methods. Plasmids are available upon requests.

## AUTHOR CONTRIBUTIONS

C.V. : Investigation, Conceptualization, Supervision, Funding acquisition, Writing – original draft ; M.B.: Investigation, Conceptualization, Supervision, Writing-review and editing ; A.R.: Investigation ; D.V.: Investigation, Conceptualization, Software, Writing-review and editing ; D. L.: Investigation ; F. M: Investigation, Writing-review and editing ; F. E.: Conceptualization, Writing-review and editing ; G.L.: Investigation ; N.H.: Investigation ; L.L.: Investigation ; M.F.: Investigation ; A. S.: Conceptualization, Investigation, Writing-review and editing ; O. B-S.: Funding acquisition ;; H.L.: Conceptualization, Writing-review and editing ; N. B.: Conceptualization, Investigation, Writing-review and editing ; P.C.: Funding acquisition, Writing-review and editing ; M.D.: Conceptualization, Writing-review and editing ; J.T.: Conceptualization, Investigation, Funding acquisition, Supervision, Writing – original draft

## ACKNOWLEDGMENTS

We thank C. Delmas from the CRCT technological facilities CRCT platform), Dr. N. Verstraete from Dr. V. Pancaldi’s team (CRCT).

