## Supplemental figures and tables for "The intrinsically disordered region of the E3 ubiquitin ligase TRIP12 induces the formation of chromatin condensates and interferes with DNA damage response"

Vargas C. et al.

#### Supplemental Figure 1

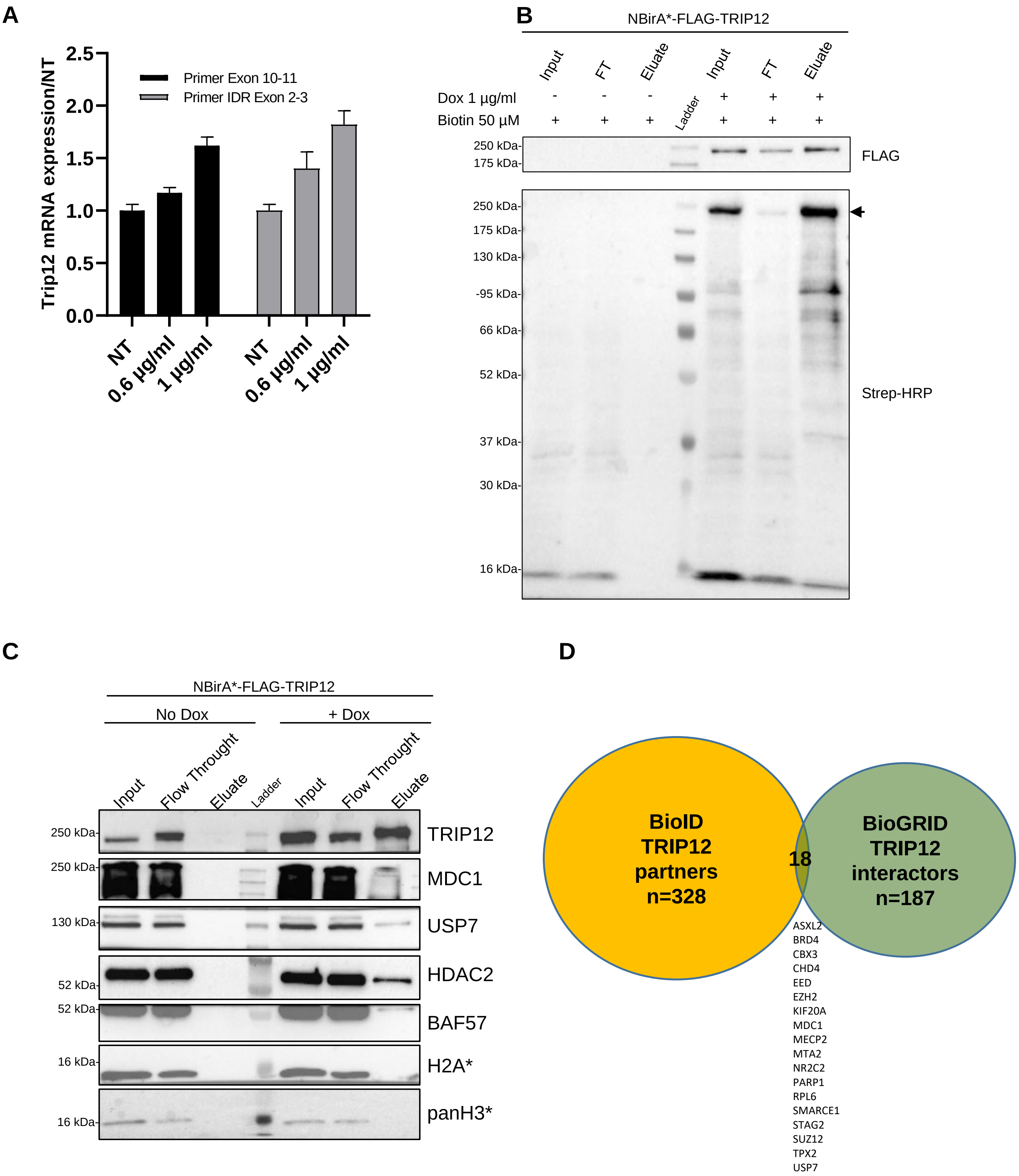

**Supplemental Figure 2**

**A**

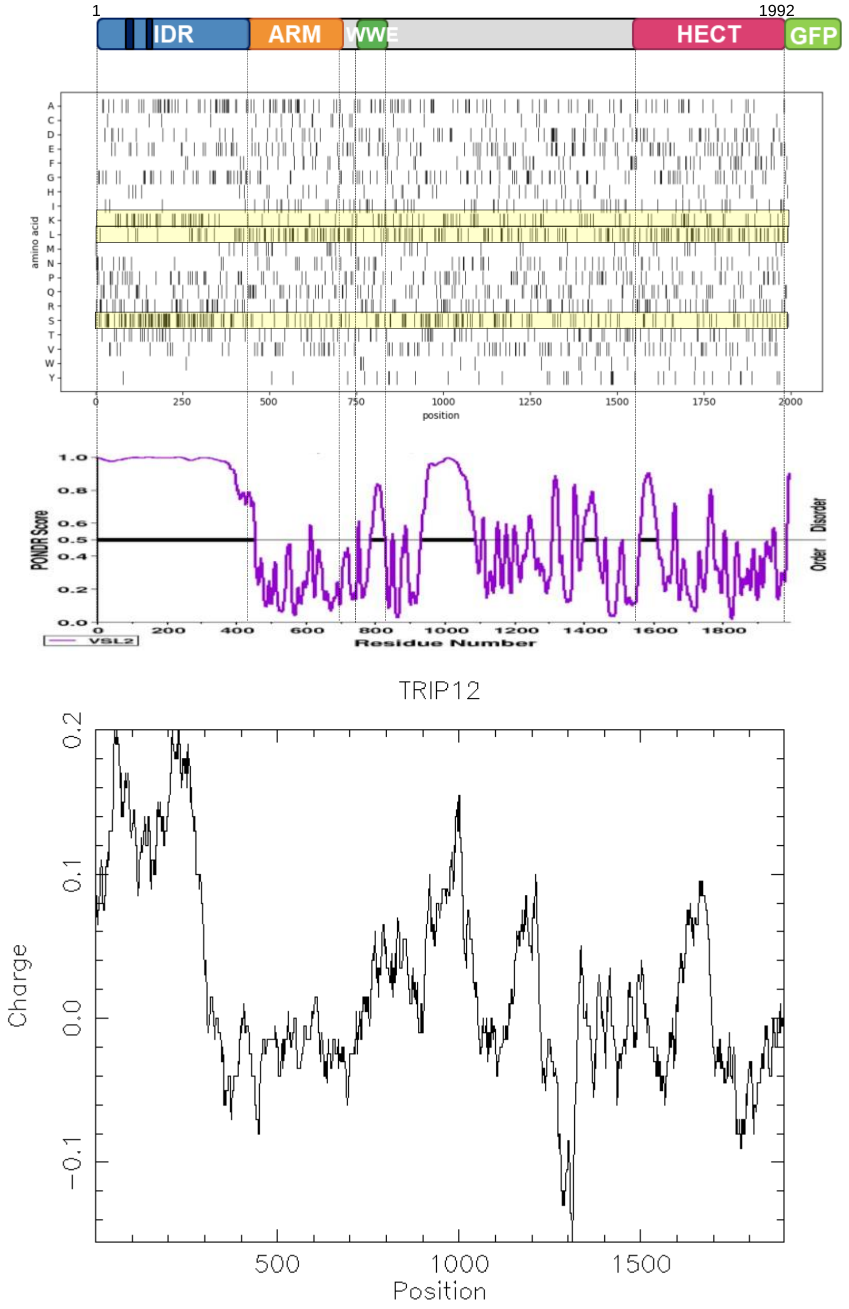


**B**

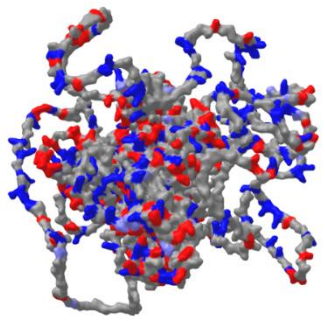

**C**

**Query protein is NULL (Q14669)**

**4 NoLSs are predicted in this protein:**

TESPSETNKPHSKSKKRHLDQEQQLKSA (between positions 98 and 125)

ATGGSRSQKRKRTESSCVKSG (between positions 140 and 160)

AKLASLRKSTKKRSESPPAE (between positions 297 and 316)

LSDVLKRKRLPKRGPRRPKYSPPRDDD (between positions 996 and 1022)

**Position in full-length protein (NoLSs shown in red):**

MSNRPNNNPGGSLRRSQRNTAGAQPQDDSIGGRSCSSSSAVIVPQPEDPDRANTSERQKT
GQVPKKDNSRGVKRSASPDYNRTNSPSSAKKPKALQHTESPSETNKPHSKSKKRHLDQEQ
QLKSAQSPSTSKAHTRKSGATGGSRSQKRKRTESSCVKSGSGSESTGAEERSAKPTKLAS
KSATSAKAGCSTITDSSSAASTSSSSSAVASASSTVPPGARVKQGKDQNKARRSRSASSP
SPRRSSREKEQSKTGGSSKFDWAARFSPKVSLPKTKLSLPGSSKSETSKPGPSGLQAKLA
SLRKSTKKRSESPPAELPSLRRSTRQKTTGSCASTSRRGSGLGKRGAAEARRQEKMADPE
SNQEAVNSSAARTDEAPQGAAGAVGMTTSGESESDDSEMGRLQALLEARGLPPHLFGPLG
PRMSQLFHRTIGSGASSKAQQLLQGLQASDESQQLQAVIEMCQLLVMGNEETLGGFPVKS
VVPALITLLQMEHNFDIMNHACRALTYMMEALPRSSAVVVDAIPVFLEKLQVIQCIDVAE
QALTALEMLSRRHSKAILQAGGLADCLLYLEFFSINAQRNALAIAANCCQSITPDEFHFV
ADSLPLLTQRLTHQDKKSVESTCLCFARLVDNFQHEENLLQQVASKDLLTNVQQLLVVTP
PILSSGMFIMVVRMFSLMCSNCPTLAVQLMKQNIAETLHFLLCGASNGSCQEQIDLVPRS
PQELYELTSLICELMPCLPKEGIFAVDTMLKKGNAQNTDGAIWQWRDDRGLWHPYNRIDS
RIIEQINEDTGTARAIQRKPNPLANSNTSGYSESKKDDARAQLMKEDPELAKSFIKTLFG
VLYEVYSSSAGPAVRHKCLRAILRIIYFADAELLKDVLKNHAVSSHIASMLSSQDLKIVV
GALQMAEILMQKLPDIFSVYFRREGVMHQVKHLAESESLLTSPPKACTNGSGSMGSTTSV
SSGTATAATHAAADLGSPSLQHSRDDSLDLSPQGRLSDVLKRKRLPKRGPRRPKYSPPRD
DDKVDNQAKSPTTTQSPKSSFLASLNPKTWGRLSTQSNSNNIEPARTAGGSGLARAASKD
TISNNREKIKGWIKEQAHKFVERYFSSENMDGSNPALNVLQRLCAATEQLNLQVDGGAEC
LVEIRSIVSESDVSSFEIQHSGFVKQLLLYLTSKSEKDAVSREIRLKRFLHVFFSSPLPG
EEPIGRVEPVGNAPLLALVHKMNNCLSQMEQFPVKVHDFPSGNGTGGSFSLNRGSQALKF
FNTHQLKCQLQRHPDCANVKQWKGGPVKIDPLALVQAIERYLVVRGYGRVREDDEDSDDD
GSDEEIDESLAAQFLNSGNVRHRLQFYIGEHLLPYNMTVYQAVRQFSIQAEDERESTDDE
SNPLGRAGIWTKTHTIWYKPVREDEESNKDCVGGKRGRAQTAPTKTSPRNAKKHDELWHD
GVCPSVSNPLEVYLIPTPPENITFEDPSLDVILLLRVLHAISRYWYYLYDNAMCKEIIPT
SEFINSKLTAKANRQLQDPLVIMTGNIPTWLTELGKTCPFFFPFDTRQMLFYVTAFDRDR
AMQRLLDTNPEINQSDSQDSRVAPRLDRKKRTVNREELLKQAESVMQDLGSSRAMLEIQY
ENEVGTGLGPTLEFYALVSQELQRADLGLWRGEEVTLSNPKGSQEGTKYIQNLQGLFALP
FGRTAKPAHIAKVKMKFRFLGKLMAKAIMDFRLVDLPLGLPFYKWMLRQETSLTSHDLFD
IDPVVARSVYHLEDIVRQKKRLEQDKSQTKESLQYALETLTMNGCSVEDLGLDFTLPGFP
NIELKKGGKDIPVTIHNLEEYLRLVIFWALNEGVSRQFDSFRDGFESVFPLSHLQYFYPE
ELDQLLCGSKADTWDAKTLMECCRPDHGYTHDSRAVKFLFEILSSFDNEQQRLFLQFVTG
SPRLPVGGFRSLNPPLTIVRKTFESTENPDDFLPSVMTCVNYLKLPDYSSIEIMREKLLI
AAREGQQSFHLS

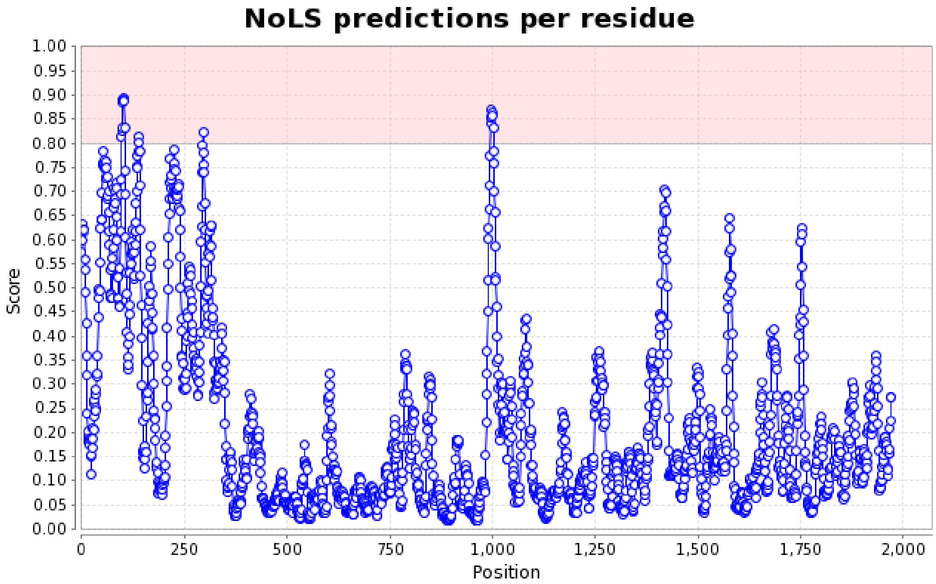

**D**

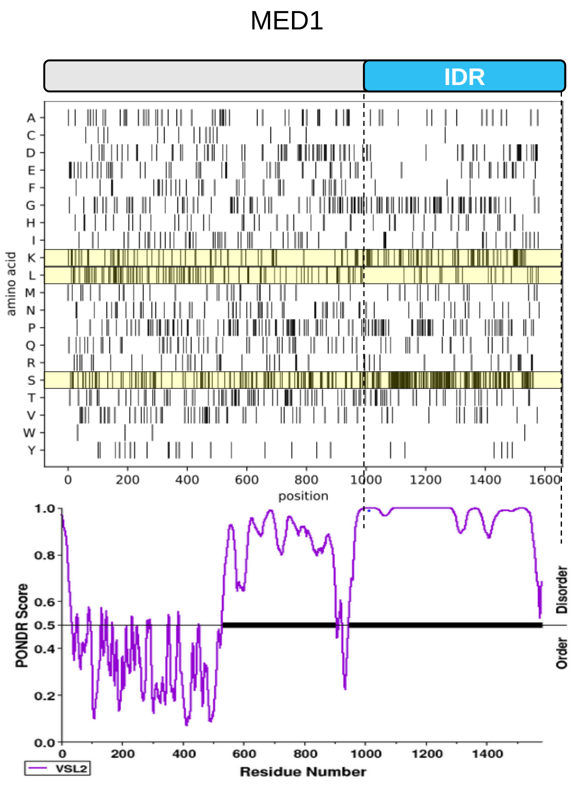
MED1
IDR
amino acid
A
C
D
E
F
G
H
I
K
L
M
N
P
Q
R
S
T
V
W
Y
0
200
400
600
800
1000
1200
1400
1600
position
PONDR Score
1.0
0.8
0.6
0.5
0.4
0.2
0.0
Disorder
Order
VSL2
Residue Number

### Supplemental Figure 3

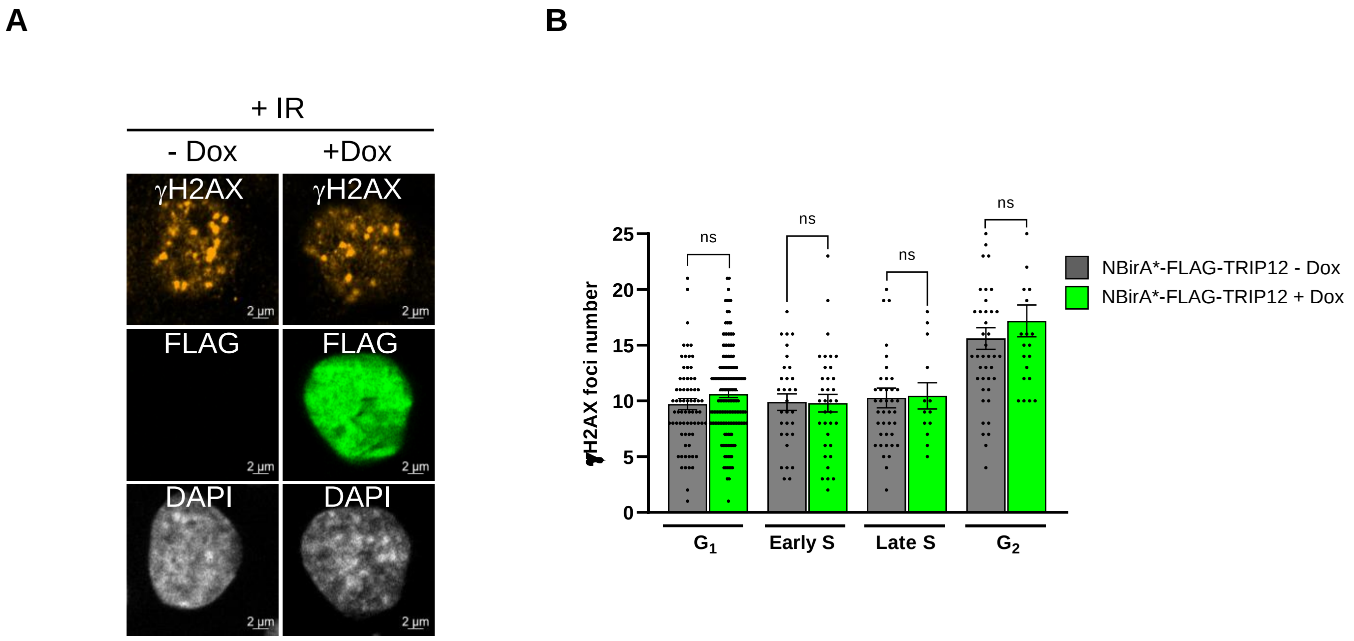

**Supplemental Table 1: List of primers**

| **qPCR** | **Sequences** |
|---|---|
| Trip12 exon 10-11 Forward | 5’-ATATTATGAACCATGCTTGTCGAG-3’ |
| Trip12 exon 10-11 Reverse | 5’-CTCTGCCACATCAATACACTGAAT-3’ |
| Trip12 exon 2-3 Forward | 5’- CTGTGCTAGTACCAGTCGGC-3’ |
| Trip12 exon 2-3 Reverse | 5’-GAGCTTCATCTGTCCGAGCA-3’ |
| Cyclophilin A Forward | 5’-GTCAACCCCACCGTGTTCTT-3 |
| Cyclophilin A Reverse | 5’-CTGCTGTCTTTGGGACCTTGT-3’ |
| Gapdh Forward | 5’-CAATGACCCCTTCATTGACC-3’ |
| Gapdh Reverse | 5’-GTACTCGCTCCTGGAAGATG-3’ |
| **Plasmid construction** | |
| MED1 NLS IDR Gateway Forward | 5’-GGGGACAAGTTTGTACAAAAAAGCAGGCTTGACCATGGCGCCGAAGA AGAAGCGAAAGGTCGAGCATCACAGTGGTAGTCAGGGT-3’ |
| MED1 NLS IDR Gateway Reverse | 5’- GGGGACCACTTTGTACAAGAAAGCTGGGTTGTGAAGATCATCTTCCTCCCC-3’ |

**Supplemental Table 2: List of antibody and conjugates**

| Proteins | Immunofluorescence dilution | Western blot dilution | RRID | References |
|---|---|---|---|---|
| TRIP12 | 1/1000 | 1/1000 | AB_1264344 | Bethyl A301-814A |
| TRIP12 | 1/1000 | NA | AB_2675340 | Sigma HPA 036835 |
| GFP | 1/500 | 1/1000 | AB_523903 | Novus Bio NB100-1770 |
| PCNA | N/A | 1/5000 | AB_628110 | Santa-Cruz SC-56 |
| 53BP1 | 1/2000 | N/A | AB_10003037 | Novus Bio NB 100-304 |
| γH2AX | 1/500 | N/A | AB_350295 | Novus Bio NB 100-384 |
| MDC1 | 1/1000 | 1/1000 | AB_10001489 | Novus Bio NB-100-395 |
| FLAG tag | 1/2000 | 1/5000 | AB_262044 | Sigma F1804 |
| FITC-Streptavidin | 1/1000 | N/A | - | eBioscience11-4317-87 |
| HSP90 | NA | 1/1000 | AB_2121214 | Cell Signaling 4874 |
| HRP-Streptavidin | NA | 1/2000 | x | ThermoFisher S911 |
| CYCLIN A2 | 1/1000 | N/A | AB_1949884 | Genetex 103042 |
| CYCLIN A2 | 1/1000 | N/A | AB_631329 | Santa Cruz SC751 |
| GAPDH | NA | 1/2000 | AB_10622025 | Cell Signaling 5174 |
| HA tag | 1/1000 | 1/1000 | AB_1549585 | Cell Signaling 3724 |
| Lamin B1 | 1/500 | N/A | AB_2136290 | Proteintech 12987-1-AP |
| CREST | 1/1000 | N/A | - | [35] |
| HP1 alpha | 1/100 | N/A | AB_2614983 | Active motif 39977 |
| H3K9me3 | 1/1000 | N/A | AB_306848 | Abcam ab8898 |
| EZH2 | 1/1000 | N/A | AB_10694683 | Cell Signaling 5246 |
| H3K27me3 | 1/1000 | N/A | AB_2616029 | Cell Signaling 9733 |
| H2AK119Ub | 1/1000 | N/A | AB_10891618 | Cell Signaling 8240 |
| USP7 | N/A | 1/1000 | AB_203276 | Bethyl A300-033A |
| HDAC2 | N/A | 1/1000 | AB_2118563 | Santa Cruz sc-7899 |
| BAF57 | 1/1000 | N/A | A300-810A-T | Bethyl A300-810A-T |
| H2A | 1/5000 | N/A | AB_2885986 | GeneTex GTX129418 |
| panH3 | 1/10.000 | N/A | AB_417398 | Upstate 07-690 |
| RNA pol II pSer2 | 1/500 | N/A | AB_2687450 | Active motif 61083 |
| RNA pol II pSer5 | 1/1000 | N/A | AB_2687451 | Active motif 61085 |
| Alexa Fluor 555 anti rabbit | 1/1000 | N/A | AB_2535849 | Invitrogen A-21428 |
| Alexa Fluor 448 anti goat | 1/1000 | N/A | AB_2534102 | Invitrogen A-11055 |
| Alexa Fluor 647 anti mouse | 1/1000 | N/A | AB_2535804 | Invitrogen A-21235 |
| Alexa Fluor 568 anti rat | 1/1000 | N/A | AB_2534121 | Invitrogen A-11077 |

**Supplemental Table 3: Most enriched proteins in BioID experiment with fold change (log2) and p value**

| Gene name | Log2(FC) | -Log10(q value 0.05) |
|---|---|---|
| H2AZ2 | 5,19 | 2,26 |
| TRIP12 | 4,69 | 2,28 |
| TAF7 | 4,60 | 4,18 |
| ZZZ3 | 4,24 | 6,98 |
| MDC1 | 4,14 | 2,53 |
| ZNF512B | 4,10 | 4,76 |
| TPX2 | 4,09 | 3,11 |
| RBBP4 | 4,04 | 2,98 |
| SCML2 | 4,04 | 3,48 |
| HEXIM1 | 3,98 | 4,21 |
| TAF5 | 3,98 | 4,62 |
| XPC | 3,94 | 3,98 |
| PHF2 | 3,89 | 3,78 |
| BAZ2A | 3,86 | 3,19 |
| ORC2 | 3,85 | 3,46 |
| CXorf56 | 3,85 | 4,39 |
| SGO2 | 3,84 | 7,96 |
| CDCA2 | 3,81 | 4,26 |
| PRPF19 | 3,79 | 4,45 |
| FLYWCH1 | 3,71 | 3,96 |
| NPM1 | 3,70 | 2,63 |
| LRWD1 | 3,70 | 3,29 |
| ORC5 | 3,67 | 2,28 |
| TCF20 | 3,62 | 3,66 |
| PPP1R10 | 3,62 | 3,80 |
| BRD7 | 3,59 | 5,58 |
| HIVEP1 | 3,58 | 3,29 |
| BEND3 | 3,57 | 4,14 |
| INTS12 | 3,57 | 4,26 |
| CHD7 | 3,53 | 5,62 |
| GLYR1 | 3,52 | 3,57 |
| BAZ1B | 3,51 | 3,02 |
| KPNA4 | 3,49 | 2,96 |
| CEBPB | 3,49 | 4,13 |
| KANSL1 | 3,49 | 4,64 |
| CDCA8 | 3,47 | 5,12 |
| ZNF280D | 3,47 | 3,93 |
| ZKSCAN4 | 3,45 | 3,44 |
| MBD2 | 3,44 | 4,49 |
| ZNF174 | 3,42 | 4,15 |
| RSF1 | 3,41 | 2,52 |
| NR2C2 | 3,39 | 4,05 |
| ZBTB33 | 3,35 | 5,23 |
| BRD2 | 3,34 | 3,60 |
| HMGXB4 | 3,32 | 6,24 |
| RNF2 | 3,30 | 4,84 |
| CUX1 | 3,30 | 5,56 |
| ATRX | 3,29 | 3,71 |
| RBBP7 | 3,28 | 2,68 |
| KPNA3 | 3,27 | 7,11 |
| SIN3A | 3,27 | 4,85 |
| NSD3 | 3,24 | 4,98 |
| BNC2 | 3,24 | 4,94 |
| PAF1 | 3,24 | 3,95 |
| USP7 | 3,24 | 4,35 |
| BAZ2B | 3,23 | 4,47 |
| SRCAP | 3,23 | 3,70 |
| AHCTF1 | 3,22 | 3,30 |
| CHD4 | 3,21 | 4,68 |
| ZNF280C | 3,21 | 3,38 |
| GPATCH4 | 3,20 | 4,33 |
| POGZ | 3,20 | 6,87 |
| CBX3 | 3,20 | 1,86 |
| TAF4 | 3,19 | 4,45 |
| SIN3B | 3,18 | 4,22 |
| TOX4 | 3,18 | 3,02 |
| TAF6L | 3,15 | 5,26 |
| MFAP1 | 3,14 | 5,33 |
| ZNF462 | 3,14 | 5,30 |
| WAPL | 3,12 | 4,98 |
| SUGP1 | 3,11 | 4,14 |
| ZNF148 | 3,10 | 4,27 |
| NSD1 | 3,09 | 6,17 |
| RSL24D1 | 3,07 | 3,11 |
| MIDEAS | 3,07 | 3,07 |
| ATF3 | 3,07 | 3,82 |
| SMARCA5 | 3,06 | 3,73 |
| DNTTIP2 | 3,06 | 3,50 |
| ZNF384 | 3,05 | 2,86 |
| AATF | 3,05 | 3,49 |
| NUMA1 | 3,05 | 4,46 |
| IWS1 | 3,03 | 2,85 |
| WIZ | 3,03 | 4,21 |
| HP1BP3 | 3,03 | 5,00 |
| ORC3 | 3,01 | 1,88 |
| DIDO1 | 2,99 | 5,56 |
| ADNP | 2,99 | 3,87 |
| LRIF1 | 2,98 | 3,42 |
| DDX52 | 2,98 | 4,14 |

**Supplemental Table 3**

| | | |
|---|---|---|
| MBD4 | 2,97 | 5,21 |
| ZNF512 | 2,97 | 5,95 |
| DHX8 | 2,97 | 6,55 |
| FIP1L1 | 2,96 | 5,33 |
| RREB1 | 2,95 | 3,82 |
| PRDM2 | 2,94 | 3,68 |
| INCENP | 2,93 | 4,22 |
| EPC2 | 2,92 | 3,57 |
| EHMT1 | 2,91 | 4,38 |
| EHMT2 | 2,91 | 4,44 |
| MECP2 | 2,90 | 2,92 |
| RECQL | 2,90 | 3,84 |
| KMT2A | 2,90 | 5,40 |
| JUNB | 2,88 | 2,83 |
| TRIR | 2,88 | 3,20 |
| PCGF1 | 2,86 | 5,48 |
| ZNF644 | 2,85 | 5,76 |
| BOD1L1 | 2,85 | 3,96 |
| EP400 | 2,84 | 5,03 |
| SUPT20H | 2,83 | 4,75 |
| ELOA | 2,82 | 3,66 |
| RPS7 | 2,82 | 4,60 |
| TOMM40 | 2,82 | 2,15 |
| ZNF131 | 2,81 | 3,49 |
| AFF4 | 2,81 | 4,29 |
| CHAMP1 | 2,81 | 5,89 |
| EZH2 | 2,81 | 5,12 |
| KIF20B | 2,81 | 5,99 |
| ZNF696 | 2,80 | 2,87 |
| CHD1 | 2,80 | 4,13 |
| ASH2L | 2,80 | 5,81 |
| NPM3 | 2,80 | 2,06 |
| TAF1 | 2,80 | 3,76 |
| PNISR | 2,79 | 5,86 |
| VIRMA | 2,78 | 4,81 |
| UTP3 | 2,77 | 3,76 |
| ZNF66 | 2,77 | 4,09 |
| PBRM1 | 2,76 | 6,02 |
| RIF1 | 2,75 | 5,08 |
| ZSCAN21 | 2,74 | 4,07 |
| PPP1CB | 2,74 | 4,27 |
| BRD4 | 2,73 | 5,09 |
| MKI67 | 2,72 | 3,76 |
| SUDS3 | 2,71 | 4,70 |
| HDAC2 | 2,71 | 5,36 |

| | | |
|---|---|---|
| BLM | 2,70 | 6,44 |
| KANSL3 | 2,70 | 3,48 |
| ATAD2 | 2,70 | 3,07 |
| SUZ12 | 2,70 | 5,78 |
| PHC2 | 2,68 | 5,30 |
| RNF169 | 2,67 | 4,43 |
| PHF12 | 2,67 | 4,17 |
| BAP18 | 2,66 | 1,49 |
| RFC1 | 2,65 | 4,83 |
| MBD3 | 2,65 | 4,45 |
| MPHOSPH10 | 2,64 | 4,26 |
| ARHGAP11A | 2,64 | 4,77 |
| DDX10 | 2,63 | 4,17 |
| GATAD2B | 2,62 | 6,18 |
| ASH1L | 2,62 | 5,09 |
| SMARCA1 | 2,60 | 4,60 |
| NOP10 | 2,60 | 2,96 |
| KIF18B | 2,59 | 4,32 |
| NSD2 | 2,59 | 7,14 |
| KNOP1 | 2,59 | 3,93 |
| RAI1 | 2,59 | 4,95 |
| TAF6 | 2,59 | 4,57 |
| H2AC4 | 2,58 | 1,70 |
| ARID2 | 2,58 | 5,34 |
| TPR | 2,58 | 3,65 |
| CBX1 | 2,58 | 2,37 |
| ZFC3H1 | 2,57 | 4,25 |
| ZNF318 | 2,56 | 3,63 |
| TRIML2 | 2,56 | 1,53 |
| CHD3 | 2,56 | 4,74 |
| RRP1B | 2,56 | 5,09 |
| MTA1 | 2,56 | 4,65 |
| H4C1 | 2,55 | 2,27 |
| CSNK2A2 | 2,54 | 5,47 |
| PPP2R2A | 2,53 | 3,28 |
| HNRNPUL1 | 2,53 | 3,23 |
| KAT7 | 2,53 | 2,80 |
| BMI1 | 2,53 | 4,40 |
| DLGAP5 | 2,52 | 4,56 |
| KPNB1 | 2,52 | 3,13 |
| BAZ1A | 2,52 | 3,48 |
| SMARCA2 | 2,52 | 4,67 |
| SMARCE1 | 2,52 | 3,42 |
| ZMYND11 | 2,52 | 2,89 |
| GPATCH1 | 2,51 | 3,27 |

#### Supplemental Table 3

| | | |
|---|---|---|
| MICOS13 | 2,51 | 3,08 |
| PARP1 | 2,50 | 2,86 |
| YEATS2 | 2,50 | 5,22 |
| CENPB | 2,50 | 2,60 |
| KAT14 | 2,50 | 6,00 |
| UHRF2 | 2,49 | 3,15 |
| ANKRD11 | 2,48 | 3,62 |
| AK3 | 2,48 | 4,98 |
| FAM207A | 2,48 | 5,10 |
| STAG2 | 2,47 | 3,64 |
| INO80 | 2,46 | 6,19 |
| NFIC | 2,46 | 3,19 |
| RLF | 2,46 | 4,47 |
| REXO4 | 2,46 | 4,11 |
| TFAP2A | 2,46 | 3,25 |
| GATAD2A | 2,46 | 4,49 |
| MSANTD2 | 2,45 | 3,55 |
| MAPKAPK2 | 2,44 | 1,80 |
| HDAC1 | 2,44 | 4,06 |
| RNPS1 | 2,43 | 2,41 |
| EPC1 | 2,42 | 5,27 |
| C5orf24 | 2,42 | 2,89 |
| BCLAF1 | 2,42 | 5,22 |
| BRIX1 | 2,41 | 3,20 |
| CBX5 | 2,41 | 2,88 |
| WDR43 | 2,40 | 3,17 |
| NFRKB | 2,40 | 3,56 |
| MLLT3 | 2,40 | 5,77 |
| ATAD2B | 2,37 | 2,90 |
| GNL2 | 2,37 | 7,21 |
| CEBPZ | 2,37 | 5,63 |
| NAT10 | 2,37 | 4,12 |
| RRP8 | 2,37 | 6,13 |
| CKAP2L | 2,36 | 4,75 |
| NFE2L1 | 2,36 | 2,96 |
| CHD8 | 2,35 | 4,55 |
| PHAX | 2,35 | 5,27 |
| RRP7A | 2,35 | 3,34 |
| GTF2E2 | 2,35 | 1,85 |
| ZBTB9 | 2,34 | 5,52 |
| ACIN1 | 2,34 | 5,82 |
| URB1 | 2,34 | 4,00 |
| DMAP1 | 2,33 | 4,88 |
| TOP2A | 2,33 | 2,65 |
| ATAD5 | 2,32 | 5,38 |
| JUN | 2,32 | 2,67 |
| PRPF3 | 2,32 | 5,04 |
| PHF10 | 2,32 | 5,81 |
| KDM2A | 2,32 | 3,31 |
| AURKB | 2,31 | 3,82 |
| PUM3 | 2,31 | 3,56 |
| TAF10 | 2,30 | 2,57 |
| EXOSC7 | 2,30 | 2,84 |
| POLR1G | 2,30 | 3,38 |
| APOBEC3B | 2,29 | 3,76 |
| EED | 2,29 | 4,64 |
| SRFBP1 | 2,29 | 5,61 |
| EMSY | 2,28 | 4,84 |
| RBM10 | 2,28 | 5,08 |
| PHRF1 | 2,28 | 4,07 |
| NOL9 | 2,28 | 5,51 |
| NBN | 2,28 | 4,09 |
| ASXL2 | 2,28 | 4,49 |
| NCL | 2,28 | 4,40 |
| PCNP | 2,28 | 2,85 |
| TASOR | 2,27 | 6,49 |
| NOP14 | 2,27 | 3,17 |
| ACTL6A | 2,26 | 4,67 |
| DHX57 | 2,26 | 2,50 |
| PHF14 | 2,26 | 2,85 |
| MGA | 2,26 | 6,24 |
| NOP9 | 2,25 | 4,04 |
| DDX55 | 2,25 | 2,85 |
| CENPF | 2,24 | 5,57 |
| UQCRFS1 | 2,24 | 2,63 |
| MACROH2A1 | 2,23 | 2,94 |
| ZC3H18 | 2,22 | 2,84 |
| INTS3 | 2,22 | 4,09 |
| RSL1D1 | 2,22 | 2,23 |
| RBM27 | 2,22 | 4,51 |
| RPRD2 | 2,21 | 5,12 |
| BRMS1L | 2,21 | 4,23 |
| MLLT1 | 2,20 | 5,24 |
| ZMYND8 | 2,20 | 3,24 |
| MYBBP1A | 2,19 | 3,42 |
| CHD2 | 2,19 | 3,24 |
| MTCH2 | 2,19 | 2,12 |
| NKTR | 2,19 | 3,37 |
| U2SURP | 2,18 | 5,88 |
| EPB41L5 | 2,18 | 3,79 |

**Supplemental Table 3**

| | | |
|---|---|---|
| SMARCA4 | 2,17 | 5,15 |
| ZNF281 | 2,17 | 4,52 |
| SGF29 | 2,17 | 4,86 |
| PHIP | 2,16 | 2,73 |
| ZMYM4 | 2,15 | 7,40 |
| CACTIN | 2,15 | 2,57 |
| ZBTB21 | 2,15 | 3,12 |
| RRP36 | 2,15 | 3,20 |
| RPL6 | 2,14 | 4,68 |
| GTF3C1 | 2,14 | 5,39 |
| GTF3C3 | 2,14 | 4,64 |
| CDCA5 | 2,14 | 4,12 |
| ZFHX3 | 2,14 | 2,90 |
| CENPC | 2,14 | 3,66 |
| FNBP4 | 2,13 | 2,67 |
| SART3 | 2,13 | 3,90 |
| MEF2D | 2,13 | 1,91 |
| HCFC1 | 2,13 | 5,46 |
| ESF1 | 2,12 | 3,64 |
| ZNF770 | 2,12 | 2,68 |
| ZMYM3 | 2,12 | 3,02 |
| IPO7 | 2,11 | 4,33 |
| DDX46 | 2,11 | 3,63 |
| BCORL1 | 2,11 | 3,11 |
| KIF20A | 2,11 | 3,77 |
| KANSL2 | 2,10 | 3,45 |
| CTR9 | 2,10 | 3,70 |
| MTA2 | 2,09 | 4,12 |
| PES1 | 2,09 | 4,12 |
| ZMYM1 | 2,08 | 2,27 |
| MRGBP | 2,08 | 3,86 |
| ZNF146 | 2,08 | 1,53 |
| WRNIP1 | 2,08 | 3,92 |
| PNN | 2,08 | 5,11 |
| SPEN | 2,07 | 3,35 |
| MGST3 | 2,07 | 2,42 |
| POLR3C | 2,07 | 2,62 |
| BRCA1 | 2,07 | 4,74 |
| CBX8 | 2,06 | 4,96 |
| ZNF592 | 2,06 | 5,04 |
| TOP2B | 2,06 | 2,66 |
| TENT4B | 2,06 | 3,39 |
| ELL2 | 2,06 | 3,12 |
| KDM5A | 2,05 | 4,32 |
| PSIP1 | 2,05 | 2,57 |
| TNPO3 | 2,05 | 3,33 |
| SYF2 | 2,05 | 1,81 |
| SNRPE | 2,05 | 2,31 |
| ZC3H14 | 2,04 | 3,95 |
| SMARCC2 | 2,04 | 4,55 |
| BPTF | 2,04 | 4,07 |
| KIF22 | 2,04 | 4,16 |
| SLC25A6 | 2,03 | 1,47 |
| HIRA | 2,03 | 3,54 |
| NIPBL | 2,03 | 5,13 |
| USP36 | 2,02 | 5,19 |
| UBTF | 2,02 | 3,22 |
| LARP7 | 2,02 | 3,24 |
| TRIM26 | 2,02 | 2,91 |
| PAXBP1 | 2,01 | 3,07 |
| THRAP3 | 2,01 | 5,40 |
| HIGD1A | 2,00 | 1,82 |

#### SUPPLEMENTAL FIGURE LEGENDS

##### Supplemental figure legend 1:

**A-** Trip12 mRNA expression level in response to doxycycline treatment in NBirA*-FLAG-TRIP12 expressing HelaS3 cells determined by RT-qPCR. Cells were treated or not with doxycycline at the indicated concentrations for 24 h. Trip12 mRNA expression was measured using two different sets of primers. Results are expressed as mean ±SEM obtained from two independent experiments.

**B-** Biotinylated protein pull-down obtained from NBirA*-FLAG-TRIP12 expressing HelaS3 cells. Cells were treated or not with doxycycline (1 µg/ml) for 72 h and with biotin (50 µM) for 24 h. Biotinylated proteins were visualized in input, flow-through (FT) and eluate fractions by Western blot using streptavidin-HRP conjugate. NBirA*-FLAG-TRIP12 expression was visualized using an anti-FLAG antibody. The black arrow indicates the biotinylated form of NBirA*-FLAG-TRIP12.

**C-** Validation of indicated protein streptavidin pull-down in input, flow-through (FT) and eluate fractions by Western blot. NBirA*-FLAG-TRIP12 expressing HelaS3 cells were treated or not with doxycycline (1 µg/ml) for 72 h and with biotin (50 µM) for 24h. * H2A and panH3 were not enriched in TRIP12 proxisome and were used as negative control.

**D-** Venn diagram of BioID TRIP12 partners (n=328) and BioGRID TRIP12 interactors (n=187). Protein in common (n=18) are listed under the diagram.

##### Supplemental figure legend 2:

**A-** Graphical representation of human TRIP12 protein sequence (top). Amino acid composition of TRIP12 protein (middle). Lysine (K), leucine (L) and serine (S) residues are highlighted in yellow (second from the top). Prediction of natural disordered regions in TRIP12 protein using PONDR® website (third from the top). Electric charge of TRIP12 protein using Expasy-Compute pI/MW tool (bottom).

**B-** 3D representation of positive (blue) and negative (red) electric patches on TRIP12 protein using AlphaFold.

**C-** Bio-informatic prediction of nuleolar localization signals in TRIP12 protein (#Q11469) sequence determined by Nucleolar localization sequence Detector software.

**D-** Graphical representation of human MED1 protein sequence (top). Amino acid composition of MED1 protein (middle). Lysine (K), leucine (L) and serine (S) residues are highlighted in yellow. Prediction of natural disordered regions in MED1 protein using PONDR® website (bottom).

##### Supplemental figure legend 3:

**A-** Representative images of γH2AX foci, and FLAG in irradiated NBirA*-FLAG-TRIP12 expressing HelaS3 cells treated or not with doxycycline (1 µg/ml for 48 h). Cells were X ray-irradiated (1 Gy) 1 h before the end of doxycycline treatment. The determination of the cell cycle phase is described in Materials and Methods. Nuclei were counterstained with DAPI.

**B-** Quantification of γH2AX foci in irradiated NBirA*-FLAG-TRIP12 expressing HelaS3 cells treated or not with doxycycline during the different phase of the cell cycle determined by confocal microscopy and FIJI macro command. Each dot corresponds to one cell. Results are expressed as mean ± SEM obtained from independent.
